# Proximity labeling proteomics reveals Kv1.3 potassium channel immune interactors in microglia

**DOI:** 10.1101/2024.01.29.577122

**Authors:** Christine A Bowen, Hai M Nguyen, Young Lin, Pritha Bagchi, Aditya Natu, Claudia Espinosa-Garcia, Erica Werner, Prateek Kumar, Brendan R Tobin, Levi Wood, Victor Faundez, Heike Wulff, Nicholas T Seyfried, Srikant Rangaraju

## Abstract

Microglia are the resident immune cells of the brain and regulate the brain’s inflammatory state. In neurodegenerative diseases, microglia transition from a homeostatic state to a state referred to as disease associated microglia (DAM). DAM express higher levels of proinflammatory signaling, like STAT1 and TLR2, and show transitions in mitochondrial activity toward a more glycolytic response. Inhibition of Kv1.3 decreases the proinflammatory signature of DAM, though how Kv1.3 influences the response is unknown. Our goal was to establish the potential proteins interacting with Kv1.3 during the TLR4-mendiated transition to DAM. We utilized TurboID, a biotin ligase, fused to Kv1.3 to evaluate the potential interacting proteins with Kv1.3 via mass spectrometry in BV-2 microglia during an immune response. Electrophysiology, western blots, and flow cytometry were used to evaluate Kv1.3 channel presence and TurboID biotinylation activity. We hypothesized that Kv1.3 contains domain-specific interactors that vary during an TLR4-induced inflammatory response, some of which are dependent on the PDZ-binding domain on the C-terminus. We determined that the N-terminus of Kv1.3 is responsible for trafficking Kv1.3 to the cell surface and mitochondria (*e.g.* NUNDC, TIMM50). The C-terminus interacts with immune signaling proteins in an LPS-induced inflammatory response (*e.g.* STAT1, TLR2, and C3). There are 70 proteins that rely on the c-terminal PDZ-binding domain to interact with Kv1.3 (*i.e.* ND3, Snx3, and Sun1). Overall, we highlight that the Kv1.3 potassium channel functions beyond outward flux of potassium in an inflammatory context and contributes to activity of key immune signaling proteins, such as STAT1 and C3.

**MAIN POINTS:** Kv1.3 channels are highly abundant in pro-inflammatory microglia in neurological diseases. Kv1.3 channels may regulate microglial functions by interacting with other proteins via its N and C terminal domains.

Using proximity-based proteomics, we identified several novel proteins that interact with the N and C terminus of Kv1.3 channels, some of which are domain-specific.

Kv1.3 channels in microglia interact with many immune signaling proteins, including Tlr2, Stat1 and integrins.

Under homeostatic conditions, the N-terminus of Kv1.3 interacts with proteins involved in protein trafficking, to the cell surface and mitochondria. The PDZ-binding region was an important determinant of the C terminal interactome.

During an LPS-induced inflammatory response, the C-terminus of Kv1.3 uniquely interacts with immune and signaling proteins of disease relevance, including STAT1

## INTRODUCTION

Microglia are the resident myeloid immune cells of the central nervous system, which are recognized to play critical roles in the pathogenesis of several neurological disorders. Microglia are involved in synaptic pruning, phagocytosis and clearance of cellular debris and protein aggregates, release of trophic and toxic factors and extracellular vesicles, which play complex roles in development and disease [1, 2]. Recent advances in transcriptomic profiling have revealed heterogeneity within microglia, where some homeostatic microglia adopt a disease-associated microglia (DAM) signature[3, 4]. DAM signatures appear to be conserved across several chronic neuroinflammatory and neurodegenerative disorders including Alzheimer’s disease (AD), Parkinson’s disease (PD), multiple sclerosis, and ischemic [5–7]. Within DAM, functional heterogeneity is also present, such that a sub-set of DAM phenotypes may have detrimental effects via increased synaptic phagocytosis and release of neurotoxic factors that have damaging effects on neurons and overall neurological function, by promoting premature cellular death and impaired pathological protein clearance [8]. Regulators of these pro-inflammatory DAM functions in neurological diseases, therefore, represent potential therapeutic targets for neuro-immunomodulation. The Kv1.3 potassium channel has emerged as a regulator of pro-inflammatory functions of DAM. Pharmacological blockade of Kv1.3 channels has also been shown to reduce neuropathology in mouse models of AD, PD and stroke models [5, 9–11]. However, the molecular mechanisms that allow Kv1.3 channels to regulate immune functions of microglia, are not fully understood.

The *Kcna3* gene encodes the Kv1.3 potassium channel protein, which homo-tetramerizes to form functional voltage-activated outward-rectifying K+ channels [12, 13]. K+ efflux via Kv1.3 fine tunes membrane potential following membrane depolarization, which in turn regulates calcium flux in immune cells such as effector memory T cells and microglia [14]. Beyond the cell surface, there is also biophysical evidence for the presence of Kv1.3 channels in the inner mitochondrial membrane, which may allow intracellular Kv1.3 channels to regulate metabolism, apoptosis and immune functions [15, 16]. While Kv1.3 channels can regulate membrane potential and calcium flux, Kv1.3 channels may also regulate immune pathways via direct protein-protein interactions at the plasma membrane [9, 17]. Based on the observed co-localization of Kv1.3 channels with integrins, receptors and immune signaling proteins in T cells, there is a strong possibility that Kv1.3 also regulates immune signaling through directly interacting with proteins associated with immune function[18, 19].

Each Kv1.3 monomeric sub-unit contains six transmembrane domains, with both the N and C termini facing the cytosol [12, 20, 21]. The N-terminus of Kv1.3 contains domains responsible for channel tetramerization, sub-unit assembly and localization to the membrane. The C-terminus contains a PDZ-binding domain, which can interact with PSD-95 and signaling proteins which form scaffolds for ERK signaling [17, 22]. Removal of the PDZ-binding domain results in changes of the Kv1.3 function and localization [23]. The Kv1.3 channel also interacts with a homotetramer of Kvβ proteins (Kvβ1-3), which further extends the breadth of potential interactors of Kv1.3 channels [22]. Many mechanistic aspects of Kv1.3 channel interactors and function have been investigated in T cells. However, the protein interactions with the N and C termini of Kv1.3 channels in microglia are less well characterized. Given the molecular and functional heterogeneity in microglial responses within DAM, it is also possible that the protein-protein interactome of Kv1.3 channels may also vary, based on the activation context of microglia.

To test this hypothesis, we investigated the N and C terminal interactomes of Kv1.3 channels in microglia, and their relationship to the activation state of microglia. We hypothesized that Kv1.3 contains domain-specific interactors, some of which are dependent on the PDZ binding domain on the C-terminus, and shift toward immune function annotated proteins during an LPS-response in microglia. To accomplish this, we utilized proximity-dependent biotinylation of Kv1.3 by fusing the biotin ligase TurboID to the N or C terminus of Kv1.3 in HEK-293 and BV-2 microglial cell lines [24]. Utilizing the BV-2 microglia cell line is more efficient than primary microglia for stable cell line transduction and generating sufficient material for proteomics studies [25]. Biotinylated proteins were then enriched and quantified using mass spectrometry-based proteomics. After verifying that these TurboID-Kv1.3 fusions did not impact channel localization and function, we identified N and C terminal protein-protein interactomes, highlighting several novel domain-specific associations between Kv1.3 channels and immune function. We also imposed an inflammatory challenge with lipopolysaccharide (LPS) to induce a pro-inflammatory microglial state which mimics the TLR activation that occur in in AD and PD[26, 27]. We assessed the relationship between microglial pro-inflammatory activation, and the Kv1.3 channel interactomes. Our studies identified several novel domain-specific immune protein interactions of Kv1.3 channels in microglia, including STAT1 and C3 at the C-terminus and TIMM50 at the N-terminus, which may explain how microglial Kv1.3 channels participate in regulating diverse immune mechanisms of neurological diseases.

## METHODS

### PLASMID and LV design

Five plasmid constructs were designed through the Emory University custom cloning core using cloning strategies summarized in Table 1.

**Table 1.**
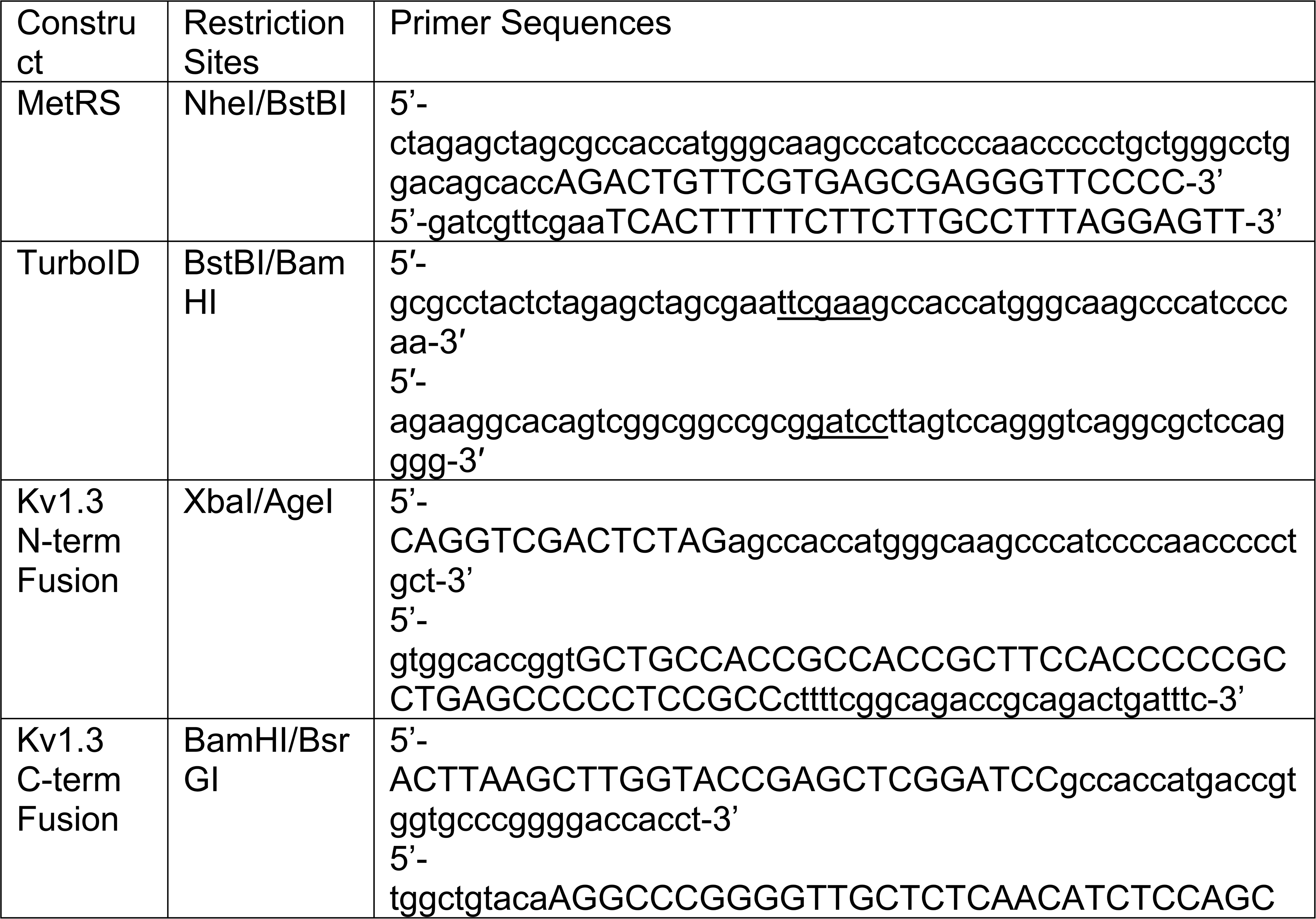
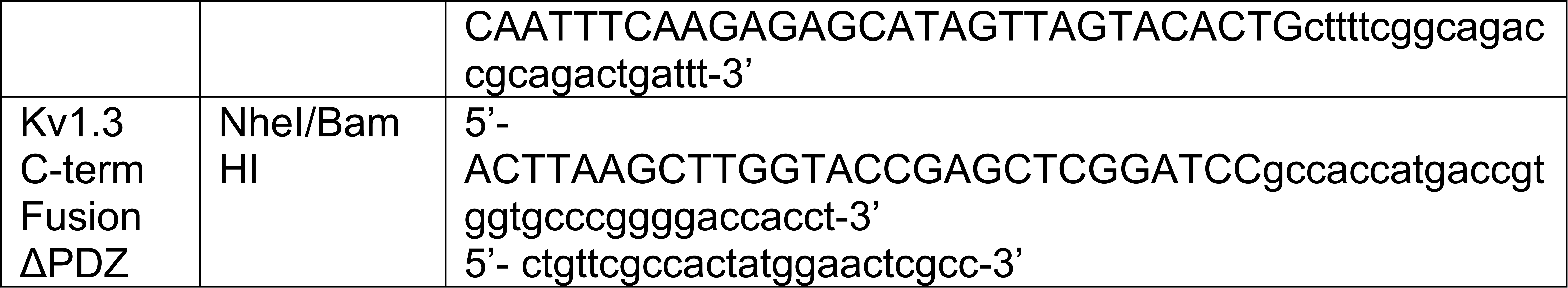
Constructs Designed for experiments conducted.

The TurboID construct was previously described in Sunna et al. [28]. The V5-TurboID-NES plasmid (Addgene, #107169) was transformed using a competent E. coli strain DH5α according to the manufacturer protocols. QIAfilter Plasmid kits (Midi prep kit; Qiagen; catalog no.: 12243) were utilized to purify plasmid DNAs following the manufacturer’s protocol. Restriction sites were introduced via the PCR primers and the V5-TurboID-NES sequence was subcloned into pCDH-EF1-MCS-BGH-PGK-GFP-T2A-Puro (CD550A-1) and sequenced (**Table 1**). This construct was utilized as a positive control for biotinylation of proteins.

The construct utilized for a transfection control comprised of an overexpression of methionine-tRNA synthetase (MeTRS) (AddGene, pMarsL274G) were created and sequenced using protocol described above.

Each of the *Kcna3* based constructs was on a FUGW backbone with *Kcna3* isolated from cDNA with a GS rich 15 amino acid linker (5’-GGCGGAGGGGGCTCA-3’x3), a v5 tag, and TurboID either on the 3’ or 5’ end of the *Kcna3* gene. These constructs were formed via protocols described in Sunna et *al.* [29].

The Kv1.3 C-term FusionΔPDZ construct had a truncated Kcna3. The deletion of TDV, which are the last 3 amino acids are described as essential to Kv1.3 function and localization [23].This truncated Kcna3 was fused with the same 15 amino acid linker, a v5 tag, and TurboID. Puromycin resistance was used as a selection marker at a separate location of the plasmid. DNA sequencing was used to confirm plasmid orientation and correct insertion.

Plasmids were packaged into lentivirus (LV) by the Emory University Viral Vector Core and purified as described below. HEK-293FT (Invitrogen) cells were maintained in complete medium (4.5g/L Glucose and L-Glutamine containing DMEM supplemented with 10% FBS and 1% Pen-Strep) and incubated at 37 °C, 5% CO_2_. One day before transfection, HEK-293FT cells were seeded onto five 150mm plates at a density of 1x10^7^ cells per plate in 20mL of complete medium. The cells were approximately 70-80% confluent at the time of transfection. The day of transfection, the DNA mixture prepared as the following: 53μg of lentiviral plasmid, 35μg of pCMVΔ 8.9 and 17.5μg of pVSVG in 4.5mL of ddH2O, 0.5mL of 1.5M NaCl. The polyethyenimine (PEI) mixture prepared as the following: 0.84mL of 7.5mM PEI, 0.5mL of 1.5M NaCl in 3.66mL of ddH_2_O. The DNA mixture and PEI mixture were then combined. This 10mL solution was vortexed 20sec and incubated for 20min at room temperature. Then 2mL of the mixture was added drop wise to each dish then incubated for 48h before harvesting.

For plasmid purification, the supernatants (media) containing lentivirus were collected 48h and 72h post-transfection, combined and then centrifuged at 500 x g for 5min at 4 °C, followed by passage through a 0.45μm low protein binding filter. The total 200mL of supernatant was centrifuged at 28,000rpm for 2h at 40 °C in a 45Ti rotor (Beckman), which can sustain faster speeds. The virus pellets were resuspended in 500μL of PBS, incubated on ice for 30min, resuspended virus particles were combined and loaded into a 12mL of SW 41 tube, 3mL of 20% sucrose cushion, and centrifuged at 28,000rpm for 2h at 4 °C in a SW 41 rotor (Beckman). The virus pellet was resuspended in 60μL of PBS, then stored at -80 °C.

For transduction of BV-2 cells, purified virus was added to cells at an multiplicity of infection(MOI) of 10 along with 8μg/mL polybrene for 24h. Remaining LV media was then removed and cells were allowed to grow in DMEM-F12 (10% FBS, 1% Penicillin/streptomycin) for 5days. Puromycin was then added at 2μg/μL for 7days. Presence of *Kcna3* gene was confirmed with qPCR.

### Cell Culture and Maintenance

Mycoplasma free HEK-293 cells (obtained from Seyfried Lab) were grown in DMEM-F12 (10% FBS, 1% Penicillin/streptomycin) and plated at 1 million cells in a 10cm dish for protein lysis and 100,000 cells/well on Poly-L-lysine coated coverslips (coating protocol followed via manufacturer’s procedure) in 24 well plates for immunofluorescence. Mycoplasma free BV-2 cells (Obtained from Tansey lab) were grown in DMEM-F12 (10% FBS, 1% Pen/strep) at 1 million cells in a 10cm dish. Cells were allowed to adhere to the plate for 24h prior to experiments.

### Dosing and Immune Stimulation

HEK-293 cells were transfected with plasmids using the JETPRIME transfection reagent for 24h according to the manufacture’s protocol. For biotinlyation, HEK-293 cells underwent a full media change with the addition of 200μM Biotin in DMEM-F12 (10%FBS, 1%P/S) for 24h.

BV-2 cells were treated with either 100ng/mL LPS or PBS for 24h. At 23h of LPS or PBS incubation, 200μM Biotin was added in without a media change for 1h. In contrast to 24h of biotinylation used in HEK-293 studies, we limited the biotinylation duration to 1h, to increase the stringency of the interactomes of Kv1.3.

### Electrophysiology

Electrophysiological experiments were conducted on transfected HEK-293 and transduced BV-2 cells that were plated on poly-Lysine-coated glass coverslips. The cells were allowed to attach for 10min at 37 °C before starting the electrophysiological measurements using the whole-cell configuration of the patch-clamp technique at room temperature with an EPC-10 HEKA amplifier. Transfected HEK-293 cells were visualized by epifluorescence microscopy of green fluorescent protein by eEGFP-C1 plasmid cotransfection (addgene, 2487). The cells were incubated at 37°C for 10min before starting the electrophysiological measurements using the whole-cell configuration of the patch-clamp technique at room temperature with an EPC-10 HEKA amplifier. The external normal Ringer solution used contained 160mM NaCl, 4.5mM KCl, 2mM CaCl_2_, 1mM MgCl_2_, 10mM HEPES, pH 7.4, and had an osmolarity of 300mOsm. Patch pipettes were made from soda lime glass (micro-hematocrit tubes, Kimble Chase, Rochester, NY) and had resistances of 2-3 MΩ when submerged in the bath solution. The pipettes were filled with an internal solution containing 145mM KF, 1mM HEPES, 10mM EGTA, 2mM MgCl_2_, pH 7.2, 300mOsm. Series resistance and whole-cell capacitance compensation were used as criteria for electrophysiology. Current amplitudes were recorded in voltage-clamp mode and elicited using a 200-ms voltage step from -80 to +40 mV at a frequency of 0.1 Hz. Use-dependency was determined using the same protocol as above but with a pulse frequency of 1 Hz for 10 pulses. The fractional current of the last pulse was normalized to the first pulse to determine the extent of cumulative (use-dependent) inactivation. Whole-cell patch-clamp data are presented as mean ± S.D., and statistical significance was determined using a paired Student’s t-test for direct comparison between WT and TurboID-fusion constructs.

The voltage dependence of activation was examined using a step protocol where cells were depolarized for 200ms from a holding potential of -80mV to a range of potentials from -80 to +40 mV in 10-mV increments, with an interpulse duration of 30sec. The peak currents were normalized to the maximum peak current and plotted against the voltage. The reversal potential was calculated by individually fitting the resulting I-V curve of each data set with the equation: I = z * (V – Vr), where I is the normalized current amplitude, z is the apparent gating charge, V is the potential of the given pulse, and Vr is the reversal potential. Conductance was then directly calculated using the equation: G = I/(V – Vr), where G is conductance and I, V, and Vr are as described above. The conductance values were fit with the two-state Boltzmann equation: G = [1 + exp(−0.03937 × z × (V – V_1/2_))]−1, where z is the apparent gating charge and V is the potential of the given pulse, and V_1/2_ is the potential for half-maximal activation.

The activation and inactivation kinetics were examined using currents elicited by the same step protocol used for determining the current amplitude at +40 mV above. The activation and inactivation time constants were calculated using the Chebyshev method to fit the activating phase and inactivating phase, respectively, of each trace with a single exponential equation: I = A × exp[–(t – K)/τ] + C, where I is the current, A represents the relative proportions of current activating with the time constants τ, K is the time shift, and C is the steady-state asymptote. Only the constants τ are reported.

### Quantitative Real Time Polymerase Chain Reaction (qRT-PCR)

#### RNA extraction

BV-2 cells were washed 2x with cold 1x PBS, then incubated in Trizol for 5 min. A chloroform/phenol extraction was performed according to manufacturer’s protocol. RNA was suspended in 50μL of DPEC treated water. Purity and quantity of RNA was evaluated using a Nanodrop 2000 (Thermo scientific). A260/280 above 1.3 and A260/230 ≥ 2.0 were considered acceptable purity levels for RNA.

#### cDNA Synthesis

2μg of RNA was mixed with 5μL 10x RT buffer, 2.5μL Multiscribe reverse transcriptase, 2μL dNTP mix, 5μL 10x RT random Primers, and DPEC H_2_O for a 50μL reaction and incubated in an Applied Biosystems 2720 Thermocycler according to Applied Biosystems Protocol. cDNA was diluted 1:10 prior to qRT-PCR.

#### Real-Time Quantitative Reverse Transcription PCR (qRT-PCR)

A 20μL reaction was utilized with 10μLTaqman Universal Master Mix II, 1μL designated primer (listed in Table 2), 4uL DPEC H_2_O, and 5μL cDNA (diluted 1:10) according to the Applied Biosystems protocol in the MicroAmp Fast-96-well Reaction Plate sealed with MicroAmp Optical Adhesive Film. The Applied Biosystems 7500 Fast Real-Time PCR System was utilized. cDNA was probed with either *Kcna3* or the housekeeping gene *Gapdh*. Fold Change was calculated on CT values and normalized to housekeeping genes and negative control. Statistical significance was calculated based on unpaired t-test and graphed on PRISM(Version 10.1.0).

**Table 2.** Reagents List.

| **Reagent** | **Manufacturer** | **Catalog #** |
| --- | --- | --- |
| Dulbecco's Modified Eagle Medium (DMEM-F12) | Gibco | 11965-092 |
| Penicillin-Streptomycin | Gibco | 15140-122 |
| Fetal Bovine Serum (FBS) | Gibco | 26140-079 |
| JETPRIME | Polyplus transfection | 101000046 |
| Lipopolysaccharide (LPS) | Sigma-Aldrich | L4391, 1 mg |
| Polybrene | Sigma-Aldrich | S2667 |
| Puromycin | Gibco | A11138-02 |
| Recombinant mouse INF-gamma | R&D Systems | 485-MI-100 |
| 0.05% Trypsin-EDTA | Gibco | 253000054 |
| Biotin | Sigma-Aldrich | B4639-100mg |
| Trizol | Invitrogen | 15596018 |
| 10x RT buffer | Applied Biosystems | 4319981 |
| Multiscribe reverse transcriptase | Applied Biosystems | 4308228 |
| dNTP mix | Applied Biosystems | 4367381 |
| 10x RT random Primers | Applied Biosystems | 4319979 |
| Taqman Universal Master Mix II, no UNG | Applied Biosystems | 4440040 |
| MicroAmp Fast-96-well Reaction Plate | Applied Biosystems | 4346907 |
| MicroAmp Optical Adhesive Film | Applied Biosystems | 4311971 |
| *Kcna3* qPCR primer | Applied Biosystems | Mm00434599_s1 |
| *Gapdh* qPCR primer | Applied Biosystems | Mm99999915_g1 |
| Rabbit Anti-V5 tag | Abcam | ab206566 |
| Donkey anti-Rabbit, FITC | Invitrogen | A16030 |
| Streptavidin-594 antibody | Invitrogen | 21842 |
| DAPI | Sigma-Aldrich | 10236276001 |
| Prolong Diamond | Invitrogen | p36970 |
| 1x fixation buffer | eBioscience | 00-8222-49 |
| 1x permeabilization buffer | eBioscience | 00-8333-56 |
| Streptavidin, Alexa-Fluor 488 conjugate | Invitrogen | 21832 |
| Unstained OneComp ebeads compensation beads | eBioscience | 01-111-42 |
| HALT protease & phosphatase inhibitor cocktail | Thermofisher | 1861284 |
| BCA colorimetric assay | Thermofisher | 23227 |
| Bovine Serum Albumin Standards | Thermofisher | 23208 |
| 4x Laemmli buffer | Bio-Rad | 1610747 |
| BOLT 4%-12% Bis-Tris Gels | Invitrogen | NW04125BOX |
| 250kDa Protein Ladder | BioLabs | P7719S |
| iBLOT mini stack system | Invitrogen | IB23002 |
| StartingBlockT20 | Thermofisher | 37543 |
| Streptavidin, Alexa-Fluor 680 conjugate | Invitrogen | S32358 |
| Silverstain | Thermofisher | 24612 |
| Lysyl endopeptidase | Wako | 125-05061 |
| Trypsin | Promega | V5111 |
| HLB columns | Oasis | 186003908 |
| Reagent A | Thermofisher | 23222 |
| Reagent B | Thermofisher | 23224 |
| Complement proteins Luminex immunoassay | Millipore | HCMP2MAG-19K-06 |
| MAPK phosphoproteins Luminex immunoassay | Millipore | 48-660MAG |
| HSP60, 60KDa heat-shock protein | Cell Signaling | 12165 |
| β-actin | Santa Cruz Biotechnology | sc-47778 |
| IRDye800 | LI-COR | 925-32211 |
| HRP anti-mouse | Jackson ImmunoResearch | 115-035-033 |

### Immunofluorescence microscopy

Cells were washed in plate twice with 1x PBS prior to fixing with 4% PFA for 30 min at room temperature then washed three times with 1x PBS for 5min (n=2). Cells were permeabilized in 0.1%TritonX100 + 2% horse serum (HS) in 1xPBS for 30min at room temperature. Cells were incubated with rabbit anti-V5 (1:500) in 2% NHS in 1x PBS for 1h at room temperature and then washed three times with 1x PBS for 5min at room temperature. Cells were further incubated with a secondary antibody mix consisting of 1:500 Donkey anti-Rabbit FITC, and 1:500 Streptavidin-594 for 1h at room temperature followed by three 1x PBS 5min washes at room temperature. Coverslips were mounted on slides using one drop of Prolong Diamond and dried for 48h. Slides were sealed with clear nail polish for 24h and imaged using a Nikon A1R HD25 inverted confocal microscope using a 60x objective lens. Image analysis and processing was completed via ImageJ.

### Flow Cytometry

Cells (BV-2 or HEK-293) were scraped and collected using ice-cold PBS in a centrifuge tube and washed by adding extra PBS (n=3). To determine Kv1.3 presence, the cells were transferred into a flow tube and incubated with ShK-F6CA (source) [30] at a concentration of 5.5μM in 100µL PBS on ice for 30min in the dark, followed by 3 PBS washes. For each PBS washing step, 1mL of PBS was added to the flow tube and centrifuged at 2200 RPM for 3min, followed by the removal of the supernatant. The cells were kept in the dark on ice until flow cytometry was performed.

To determine biotin presence in BV-2 cells, cells were first fixed in 1x fixation buffer for 30min on ice, then washed thrice with cold 1x PBS. Cells were then permeabilized for 30min using 1x permeabilization buffer on ice. To determine the presence of biotin, fixed and permeabilized cells were incubated with Streptavidin-488 (1:500 in permeabilization buffer) and incubated for 1h on ice in the dark. After incubation, the cells were washed, as mentioned above. After the last wash, 200µL of PBS was added, vortexed and kept on ice in the dark until flow cytometry was performed. Unstained OneComp beads compensation beads and beads stained with Alexa fluoro-488, ShK-F6CA and unstained cells were used as a control. All flow cytometry data was collected on the BD Aria II instrument and analyzed using the Flow Jo software.

### Cell Lysis and protein processing

BV-2 and HEK-293 cells were rinsed twice with 1x PBS prior to scraping in PBS then spun at 800xg for 5min at RT (n=3). Cells were lysed in 8M urea in Tris-NaH_2_PO_4_ with Halt Protease inhibitor (1:100). Lysates were probe sonicated at 30% amplitude three times for 5sec on 10sec off. Lysates were centrifuged at 15000xg for 15min. Supernatants, containing solubilized proteins were processed for affinity purification (AP) and Mass Spectrometry (MS). Pellets containing not solubilized proteins were stored in -80 °C.

### Western Blot

Protein amount in each cell lysate was quantified using a BCA colorimetric assay (n=3). To confirm protein biotinylation10μg of protein were added to 4x Laemmli buffer (1:50 DTT) and boiled for 10min at 95°C and resolved on a BOLT 4%-12% Bis-Tris Gel at a current of 80mV for 15min followed by 120 mV for 40 min along with a 250kDa Ladder. Proteins were transferred to a nitrocellulose membrane using the iBLOT mini stack system. Ponceau staining for 2min were used to confirm equal loading and washed using 1x TBS for 15min. Blots were blocked for 30min using StartingBlock T20 (TBS) blocking buffer. at room temperature. Blots were then incubated with streptavidin-680 (1:10000 in Blocking Buffer) for 1h at room temperature shielded from light. Blots were washed twice in TBS-T for 10min at room temperature and twice in TBS for 5min at room temperature. Membranes were then imaged using the Odyssey Li-COR system.

### Affinity purification of biotinylated proteins

83μL of Pierce streptavidin magnetic beads (Thermo-scientific, 83817) were washed with 1mL RIPA buffer for 2min on rotation at room temperature. Protein lysates, 1mg for HEK-293 Cells and 0.5mg for BV-2, were brought up in 500μL RIPA buffer and incubated with magnetic beads on rotation at 4°C for 1h. Beads were washed twice with 1mL of RIPA buffer for 8min followed by one wash with 1mL of 1M KCl for 8min on rotation at room temperature. Beads were then washed with 1mL of 0.1M Na_2_CO_3_ for 10sec and 1mL of 2M Urea in tris-HCl buffer (pH:8) for 10sec at room temperature. For each wash, samples were spun down and incubated on magnetic rack for 2min to allow full attachment of beads to the magnet. 8μL of beads were resuspended in 30μL of 2x Laemmli buffer supplemented with 2mM biotin and 20mM DTT and boiled at 95°C for 15min. Western blot and silver stain were used to check for streptavidin labeling and protein abundance post-AP. Protein bound beads were stored at 20 °C until on-bead digestion.

### Protein Digestion and Peptide Clean Up

To prepare biotin enriched samples for mass spectrometry, protein bound streptavidin beads were washed three times with 1x PBS and then resuspended with 150µL 50 mM ammonium bicarbonate (ABC, NH_4_HCO_3_). 1mM dithiothreitol (DTT) was added to reduce samples for 30 mins on a rotor (800 rpm) at room temperature. 5mM iodoacetamide (IAA) was added to alkylate cysteines for 30min on a rotor at room temperature, protected from light. Proteins were digested overnight with 0.5µg of lysyl endopeptidase on rotation (800rpm) at room temperature. Proteins were further digested by 1µg trypsin overnight on rotation (800rpm) at room temperature. Samples were acidified to 1% Formic Acid (FA) and 0.1% trifluoracetic acid (TFA), desalted using an HLB column, and dried using cold vacuum centrifugation (SpeedVac Vacuum Concentrator).

To prepare cell lysates for mass spectrometry different concentrations were optimized for digestion. 100μg of pooled cell lysates were reduced with 1mM DTT for 30 min and alkylated with 5mM IAA for 30min protected from light. Samples were diluted two-fold in ABC and digested with 2µg of lysyl endopeptidase overnight on rotation at room temperature. Samples were further diluted to a final urea concentration of 1M and digested by 4µg trypsin overnight on rotation at room temperature. Samples were acidified to 1% Formic Acid (FA) and 0.1% trifluoracetic acid (TFA), desalted using an HLB column, and dried down using cold vacuum centrifugation (SpeedVac Vacuum Concentrator). Digestion protocol also outlined in outlined in Sunna, et al. and Rayaprolu et al. [28, 31].

### Mass Spectrometry

Derived peptides were resuspended in the loading buffer (0.1% trifluoroacetic acid, TFA) and were separated on a Water’s Charged Surface Hybrid (CSH) column (150µm internal diameter (ID) x 15cm; particle size: 1.7µm). HEK-293 and BV-2 samples were ran on same Mass Spectrometer with similar chromatography settings, only differing by number of samples run. BV-2 cell samples were run on an EVOSEP liquid chromatography system using the 15 samples per day preset gradient (88min) and were monitored on a Q-Exactive Plus Hybrid Quadrupole-Orbitrap Mass Spectrometer (ThermoFisher Scientific). Whereas, HEK-293 cell samples were run on an EVOSEP liquid chromatography system using the 30 samples per day preset gradient (44 min) and were monitored on a Q-Exactive Plus Hybrid Quadrupole-Orbitrap Mass Spectrometer (ThermoFisher Scientific). The mass spectrometer cycle was programmed to collect one full MS scan followed by 20 data dependent MS/MS scans. The MS scans (400-1600 m/z range, 3 x 10^6^ AGC target, 100ms maximum ion time) were collected at a resolution of 70,000 at m/z 200 in profile mode. The HCD MS/MS spectra (1.6 m/z isolation width, 28% collision energy, 1 x 10^5^ AGC target, 100ms maximum ion time) were acquired at a resolution of 17,500 at m/z 200. Dynamic exclusion was set to exclude previously sequenced precursor ions for 30sec. Precursor ions with +1, and +7, +8 or higher charge states were excluded from sequencing. Mass spectrometry protocols outlined in Sunna, et al. and Rayaprolu, *et al.* [28, 31].

### Protein Identification and Quantification

MS Raw files were uploaded into MaxQuant software (version 2.4.9.0), where HEK-293 data were searched against the human Uniprot 2017 database, and BV-2 data was searched against the 2020 mouse Uniprot proteome database, both of which were modified to contain target sequences for TurboID. Methionine oxidation, protein N-terminal acetylation, and deamination were variable modifications; carbamidomethyl was a fixed modification; the maximum number of modifications to a protein was 5. Label Free Quantification (LFQ) minimum ratio count was set to 1. Re-quantification, a process in Maxquant to double check peptides, was used. The minimum peptide length was set to 6 amino acids, with a maximum peptide mass of 6000Da. Identifications were matched between runs. For protein quantification, the label minimum ratio count was set to 1, peptides were quantified using unique and razor peptides. Fourier transformed MS (FTMS) match tolerance was set to 0.05Da and Ion Trap MS was set to 0.6Da. The false discovery rate (FDR) was set to 1%.

For both the enriched samples and cell lysates, Maxquant intensities were uploaded into Perseus (version 1.6.15). Categorical variables, such as potential contaminants, reverse peptides, and samples only identified by one site were all removed. Data were log_2_ transformed and filtered for missingness such that at least two samples had non-missing values within an experimental group. After this threshold, missing data were imputed based on normal distribution and matched to uniport gene names (**Supp. datasheet 1 and 2**). In enriched samples, protein groups were normalized to TurboID abundance (**Supp. Datasheet 3 and 4**). The same process was applied to all cell types and conditions described below.

### Data analysis

#### Analysis of TurboID biotinylated proteomes of HEK-293 cells

The analysis was divided into 4 groups which are “Control”, “Cterm”, “Nterm”, and “TurboID” each with a n=3 and a final dataset of 2122 proteins. The C-term fusion ΔPDZ was excluded from analysis due to inconsistent labeling. Principal component analysis (PCA) of affinity purified data was performed. Furthermore, to identify genes of interest that were differentially abundant, we performed a one-way analysis of variance (ANOVA) on all samples (n = 12), comparing the groups are “Control”, “Cterm”, “Nterm”, and “TurboID” (n = 3 each). The code for the one-way ANOVA analysis was adapted from the “parANOVA” repository on GitHub (https://github.com/edammer/parANOVA). Volcano plot comparisons were generated for the following data. The proteins which have a raw p-value of </= 0.05 and a Log_2_FC of +/-1 are considered statistically significant.

#### Analysis of TurboID biotinylated proteomes of BV-2 cells

The finalized data set of 1412 proteins were identified. A smaller number of proteins were identified in BV-2s likely just because there were less proteins biotinylated. We performed principal component analysis (PCA) for analyzing high-dimensional datasets which helped to distinguish patterns, relationships in the dataset and identify the main sources of variation.

Differential enrichment analysis (DEA) was performed. Significantly differentially abundant proteins were identified with an unadjusted p-value ≤ 0.05 (nominal p values are widely accepted for volcano analysis and hence were chosen for this analysis over BH corrected to avoid over filtering of genes and data) and a Log_2_FC of +/-1 for up and decreased proteins considered statistically significant respectively. Following the roadmap for statistical analysis, the “parANOVA” repository on GitHub (https://github.com/edammer/parANOVA) was referenced for the one-way ANOVA code and was implemented. Volcano plots were created to represent results from differential abundance analyses. Morpheus (Morpheus, https://software.broadinstitute.org/morpheus) was used to create visual heat maps of proteins abundance. Individual proteins were colored based on z-score, where the darker shades of red indicate +1 and the darker shades of blue indicates -1. Hierarchical clustering arranged proteins based on groups.

#### Analysis of whole BV-2 Cell Lysates

The input data was loaded and shared in the supplemental file. Post processing the finalized dataset identified 4152 observations or proteins. To distinguish patterns, relationships in the dataset and identify the main sources of variation, we performed principal component analysis (PCA) for analyzing high-dimensional datasets.

Differential enrichment analysis was performed. Significantly differentially enriched proteins were identified via unadjusted p-value ≤ 0.05 and fold change of +/-1. Following down the roadmap for statistical analysis, the “parANOVA” repository on GitHub (https://github.com/edammer/parANOVA) was referenced for the one-way ANOVA code and was implemented for analysis. The PBS Group was compared to the LPS Group with a n=5 of each and Volcano plot was generated for the comparison. “KCNA3” and “TurboID” were highlighted as significant proteins to clarify biological insight from the pertinent dataset.

### Gene Ontology analysis

#### HEK-293 cells

Gene set enrichment analysis (GSEA) was performed utilizing the software AltAnalyze (version 2.1.4)[32, 33]. Z-score of higher than 1.96 was used to identify significantly enriched genes. In the HEK-293 cells, the DEA comparison between N-terminal and C-terminal were used to create an input list that was unique to the N-terminus of Kv1.3 and a list of proteins enriched with the C-terminus of Kv1.3. These lists were compared to all of the proteins captured normalized to TurboID abundance. The most abundant genes were listed along with the STRING analysis.

#### BV-2 Cells

Gene set enrichment analysis (GSEA) was performed utilizing the software AltAnalyze (version 2.1.4)[32, 33]. Z-score of higher than 1.96 was used to identify significantly enriched genes The DEA comparison of both N and C termini compared to global TurboID presence was used to calculate an input list of all microglial Kv1.3 interactors. The DEA comparison between N and C termini was used to create an input list specifically for Kv1.3 N-terminal interactors. For the LPS conditions, the DEA comparison for the c-terminal fusion with PBS or LPS was used to create lists of proteins more abundant in the PBS conditioned media or LPS conditioned media. Lists were created for proteins that are enriched and depleted with the deletion of the PDZ binding domain on the C-terminus. Lists were referenced to all proteins abundant in mass spectrometry data that were normalized to TurboID abundance. The top 10 genes lists were selected based on highest z-score. The GSEA results were plotted with a bar graph of z-score and colored based on type of process using PRISM (Version 10.1.0).

### Protein-protein interaction network analysis (STRING)

#### HEK-293 Cells

Interacting networks were made using the STRING analysis software[34]. Network nodes represent proteins. Node Color represents gene term member, where dark colors indicate significant z-score in GSEA list and light color indicates not significant. Disconnected nodes were excluded from String network. Edges represent protein-protein association, where edge thickness indicates confidence, thickest line are highest confidence (0.900) Thinnest line low confidence (0.150).

#### BV-2 Cells

Interacting networks were made using the STRING analysis software[34]. Network nodes represent proteins of Kv1.3 interactors or Kv1.3 interactors overlapping with MITOCARTA. Node Color represents GSEA list. Disconnected nodes were excluded from String network. Edges represent protein-protein association, where edge thickness indicates confidence, thickest line are highest confidence (0.900) Thinnest line low confidence (0.150).

### Luminex

BV-2 cells were grown at 50,000 cells/well in 12-well plates and allowed to adhere to bottom of plate for 24h. Cells were then treated with LPS (100ng/mL) for 24h then exposed to INFγ (100ng/mL) for 60 min. Lysates were prepared in 8M as previously described above and in Sunna et al. [28].

Complement proteins and MAPK phosphoproteins were quantified using multiplexed Luminex immunoassays; these have been previously used without cross reactivity [31]. The analytes measured with the human complement panel are complement C1q, complement C3, complement C3b/iC3B, complement C4, complement factor B, and complement factor H. To determine cross-species reactivity between the human complement panel analytes and the murine cell line lysates, linear ranging was performed. Only analytes obeying a linear response were evaluated. Analytes detected with the MAPK panel are pATF2 (Thr71), pErk (Thr185/Tyr187), pHSP27 (Ser78), pJNK (Thr183/Tyr185), p-c-Jun (Ser73), pMEK1 (Ser222), pMSK1 (Ser212), p38 (Thr180/Tyr182), p53 (Ser15) and pSTAT1 (Tyr701). Standard protocols from the manufacturer were followed, with quarter volume loading of reagents to reduce excess antigen. Sample loadings were normalized to total protein via Pierce BCA. To identify samples biotinylated *in vitro*, an adapted Luminex protocol was followed as previously reported[31]. The overarching mechanism in the standard protocol is an analyte is immobilized using a magnetic bead specific to the analyte. The analytes are then tagged with specific biotinylated antibodies and biotinylation is detected with a streptavidin fluorophore; streptavidin-phycoerythrin. The adapted assay protocol utilizes the biotin-ligated proteome and thus omits the biotinylated antibody. Appropriate protein loadings for both standard and adapted assays were determined following a linear ranging to ensure signal above noise/background levels and to mitigate potential impact of the hook effect (false negatives) [31]. Average Net MFI values were utilized; negative values were imputed to zero. Statistical significance was calculated based on unpaired t-test and graphed in PRISM (Version 10.1.0).

### Preparation of Mitochondrial Enriched Fractions from BV-2 microglia cells

Method for mitochondrial enrichment was adapted from Wieckowski MR, et al. [36]. The BV-2 cell lines described previously: BV-2, TurboID, N-term fusion, C-term fusion, and the C-term fusion ΔPDZ were cultured at 5% CO_2_ in DMEM-F12 (10%FBS, 1%Penn/Strep. Untransduced cells were removed by adding 2µg/mL puromycin to the complete medium. After 5-7days of selection, all cell lines were seeded in 150mm Petri dishes and allowed to grow until 90% of confluency in complete medium., Cells were exposed to 200µM biotin for 24h before harvesting. Approximately 40-million cells were used to generate mitochondrial enriched fractions by differential centrifugation as described previously [36]. Experiments were performed in triplicates at the same time. Briefly, cells were detached by trypsinization, then washed with cold 1XPBS and pelleted at 800xg for 10min at 4°C. After aspirating PBS, pellets were resuspended in 1.6mL IB-1 buffer (225mM mannitol, 75mM sucrose, 0.1 EGTA and 30mM Tris-HCl pH 7.4) with HALT protease and phosphatase inhibitor cocktail and homogenized using a Teflon potter homogenizer with 25 strokes on ice. Total homogenates (Fraction A0) were transferred to a clean tube, and then centrifuged twice at 600xg for 5min at 4°C, pellets containing unbroken cells, nuclei, and heavy membranes were fraction A1. Supernatants were pelleted at 7,000xg for 10min at 4°C and the resulting pellets (Fraction A2) correspond to heavy membrane organelles, likely containing the largest presence of Kv1.3. Pellets were washed in 200µL 1B-2 buffer (225mM mannitol, 75mM sucrose, 30mM Tris-HCl pH 7.4) and centrifuged at 7,000xg for 10min at 4°C, resulting pellets are crude mitochondria (Fraction A3). For each fraction, pellets and homogenates were diluted in 200µL of 8M urea buffer and sonicated (5sec on–off pulses, Amplitude 30%) for 45sec, and then centrifuged at 21,130xg for 10min at 4°C. Supernatants were collected, and protein concentration was determined by BCA assay. 10μg of protein were used to verify mitochondrial enrichment (HSP60, 1:1000), TurboID fusion (anti-V5 tag, 1:250, IRDye800, 1:10,000) by western blot.

## RESULTS

### Validation of N and C terminus TurboID fusion constructs for mapping the Kv1.3 interactome in HEK-293 cells

To identify Kv1.3 interacting proteins, we employed proximity labeling via TurboID, wherein TurboID was fused to the N-terminus or C-terminus of Kv1.3 and then expressed in mammalian cells *in vitro* for proteomic labeling and mass spectrometry-based quantification of biotinylated proteins. TurboID is a biotin ligase that rapidly and promiscuously biotinylates proteins within a 10-30nm radius[24]. TurboID constructs were generated such that TurboID was fused to Kv1.3 with an intervening 15 amino acid linker, and a V5 tag. These included an N-terminal fusion of TurboID to Kv1.3, a C-terminal fusion of TurboID to Kv1.3 (**Fig. 1A and B**). A global expression of TurboID, where TurboID is expressed in the cytoplasm, was utilized as a positive control for biotinylation of the cellular proteome. Transfection with plasmid containing and unrelated v-5 tagged protein (MetRS) with an otherwise identical plasmid, was a transfection control. These constructs were inserted into plasmids and transfected into human embryonic kidney (HEK-293) cells. After 24h of transfection, cells were then exposed to biotin for 24h (**Fig. 1A**). We specifically chose HEK-293 cells based on their negligible basal levels of Kv1.3 channel expression, providing an optimal system for initial validation of the TurboID-Kv1.3 fusion constructs[35].

**Figure 1:**
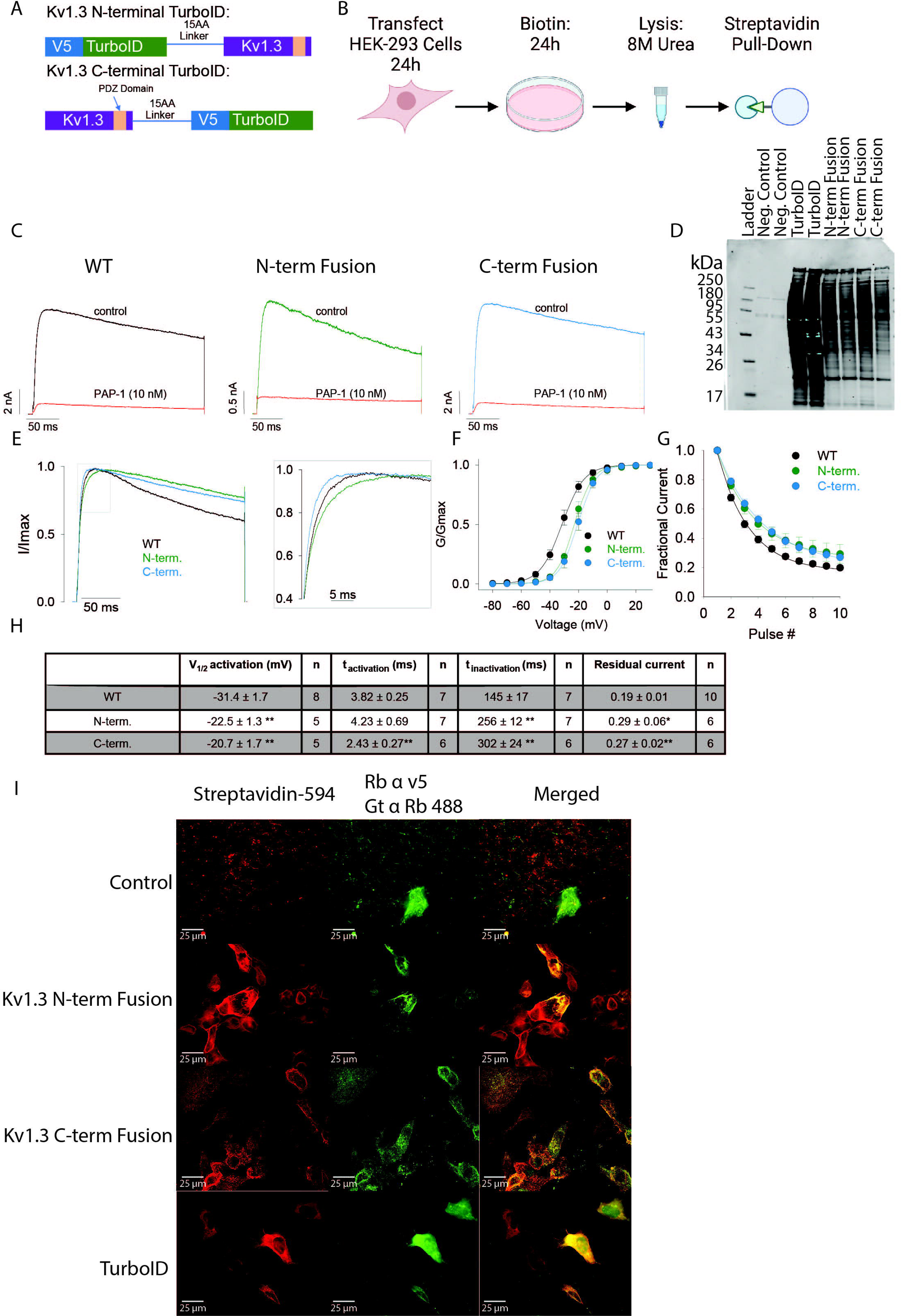
Transfected cells show Kv1.3 channel activity and biotinylation of proximal proteins. **(A)** Schematic of experimental design. HEK-293 cells were transfected with Kv1.3-turboID fusion constructs for 24h then exposed to biotin for 24h. Cells were then lysed in 8M urea and pulled-down using magnetic beads fused to streptavidin prior to mass spectrometry. **(B)** Constructs of Kv1.3 fusion with TurboID. Each fusion construct contains TurboID, a 15 amino acid linker, a V5 tag and Kv1.3. The N-terminal fusion, has TurboID located on the N-term of Kv1.3, andthe C-terminal fusion has TurboID located on the C-term of Kv1.3. . **(C)** Electrophysiology of HEK-293 cells transfected with Kv1.3-TurboID constructs show similar trace patters and respond to blockade of channel using PAP1. **(D)** Streptavidin (680) western blot shows high biotin labeling with presence of TurboID. Ponceau staining highlights consistent protein loading. **(E)** Inactivation curves for the Kv1.3 channel show that the N-terminal and C-terminal fusions have a slightly slowing inactivation rate compared to a WT control. **(F)** The conductance of the Kv1.3 channels fused to TurboID show minor shifts in the Kv1.3 channels compared to the WT control. **(G)** Electrophoresis shows the fractional current of Kv1.3 fused to TurboID is marginally higher compared to controls. **(H)** Table highlighting changes in biophysical properties with TurboID fusion to Kv1.3. **(I)** Immunofluorescence (IF) of HEK-293 cells transfected with Kv1.3-TurboID Constructs. IF highlights colocalization of biotinylated proteins (tagged with Streptavidin) with V-5 tagged TurboID. n=3*p < 0.05, **p< 0.01

We conducted electrophysiological studies on transfected HEK-293 cells using the whole-cell patch-clamp to confirm the functional expression of recombinant Kv1.3 channels at the cell surface. The current amplitude produced by the transiently transfected fusion constructs was similar to that of the transfected WT Kv1.3, indicating that the fusion with TurboID on either the N-terminus or C-terminus had no effect on trafficking and insertion of the channel into the plasma membrane. (**Fig. 1C and Supp.** Fig. 1A). At 10nM, PAP-1, a highly-selective Kv1.3 blocker, potently inhibited more than 90% of Kv1.3 currents in HEK-293 cells over-expressing WT Kv1.3 orKv1.3-TurboID fusions, confirming that all currents detected were indeed associated with transfected Kv1.3 (**Fig. 1C and Supp.** Fig. 1A). Interestingly, both constructs fused with TurboID exhibited a small shift in the voltage-dependence of activation towards more positive potentials (**Fig. 1E and 1H**), a delayed decay of use-dependent current upon repeated stimulation (**Fig. 1F and 1H**), and slower inactivation decay (**Fig. 1G and 1H**). The C-terminal fused Kv1.3 showed delayed activation kinetics compared to positive controls of HEK-293 cells transfected with unmodified Kv1.3 (**Fig. 1E and 1H**). While we did not determine the mechanism responsible for these minor changes in activation/inactivation properties of the fusion channels, we did use pharmacology to ascertain the molecular identity of Kv1.3 at the cell surface. [36]. Overall, these studies highlighted that our Kv1.3-TurboID fusion constructs are properly inserted and retained functional and pharmacological properties at the cell surface.

Immunofluorescence microscopy was performed to confirm Kv1.3 channel localization to the cell surface and inside the cell, along with colocalization with biotinylated proteins (**Fig. 1I**). Transfection control, with a non-TurboID plasmid, showed a presence of the V5 tagged MeTRS and no biotinylation (**Fig. 1I** control row). The N-terminal construct and the C-terminal construct showed similar localization of V5 and biotinylation to the cell surface, which confirmed localization of Kv1.3 and biotinylation of membrane-associated proteins that was comparable across Kv1.3 TurboID-fusion constructs. In contrast with Kv1.3-TurboID fusion constructs, we also over-expressed cytosolic TurboID (TurboID-NES not fused to any protein) [24] in HEK-293 cells, which showed expected V5 localization and biotinylation to the cytosol. These indicated that the expression of the Kv1.3 channel is likely on the plasma membrane and that the biotinylation of proteins corresponded with membrane localization of Kv1.3 channels.

Cells were then lysed in 8M Urea lysis buffer, and biotinylated proteins were affinity purified using streptavidin magnetic beads. Western blot of inputs (before streptavidin affinity purification [AP]) and after AP showed a higher level of biotinylated proteins in N-terminal and C-terminal fusions compared to negative control (**Fig 1D and Supp.** Fig. 1B and C). In contrast, the biotinylation pattern of cytosolic TurboID transfected cells was distinct from that of the Kv1. 3-TurboID fusion transfected cells (**Fig 1D and Supp.** Fig. 1B and C). These experiments established that Kv1.3 is present and functional on the surface, and that TurboID is able to biotinylate proteins in HEK-293 cells in a pattern distinct from global cytosolic TurboID expression. We proceeded with MS of biotinylated proteins to identify Kv1.3 N-terminal and C-terminal interacting proteins in HEK-293 cells.

### Kv1.3 Amino and Carboxyl terminal fusions with TurboID identifies distinct domain-interacting proteins in HEK-293 cells

After confirmation of Kv1.3 channel activity and TurboID biotinylation of proteins, we wanted to evaluate which proteins were interacting with Kv1.3. Affinity purified (AP) proteins were assessed by silver stain and Western Blot to check for quality of proteins available and efficiency of AP, a higher efficiency meaning less overall proteins in controls in the silver stain (**Supp.** Fig 1C). To evaluate Kv1.3 interactome, AP proteomes from all Kv1.3-TurboID fusion transfections were first normalized to TurboID protein abundance to account for any uneven transfection or efficiency of SA-enrichment. These normalized data were visualized using Principal component analysis (PCA). Principal Component 1 (PC1) accounted for 79% variance and clearly separated samples containing biotinylated proteins via TurboID, from non-TurboID controls (**Fig. 2A**). PC1 therefore represents proteins that were labeled by TurboID across all experimental conditions.

**Figure 2:**
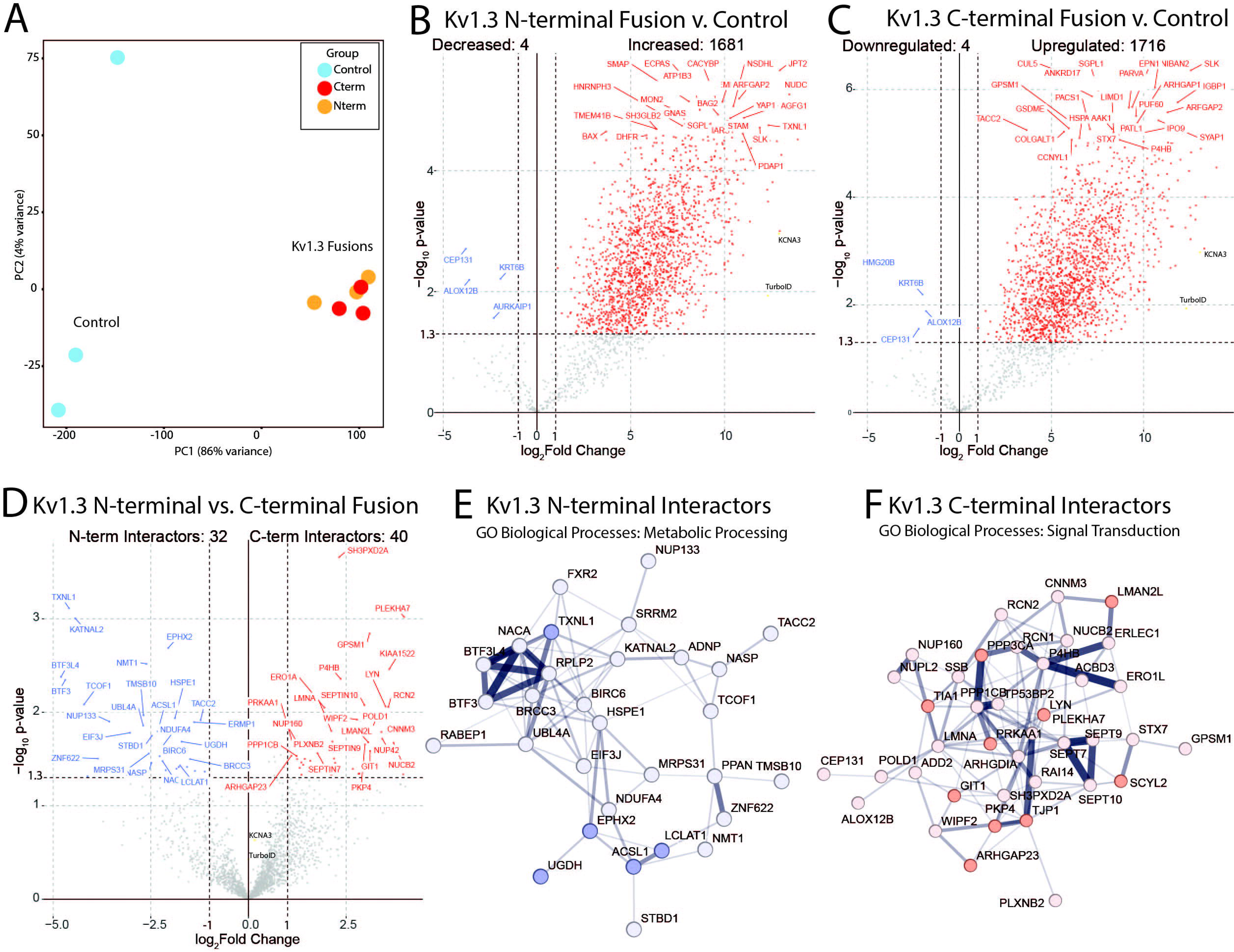
Kv1.3 interactors in HEK-293 Cells show that the N-terminus is associated with protein processing and the C-terminus is involved in signaling. **(A)** Principal Component Analysis (PCA) of mass spectrometry of biotinylated proteins shows distinct separation between control and Kv1.3-TurboID transfected cells. **(B)** Differential Abundance Analysis of N-terminal interactors shows about 1600 protein interacting with Kv1.3. **(C)** Differential Abundance Analysis of C-terminal interactors shows about 1700 proteins interacting with Kv1.3. **(D)** Differential Abundance Comparison between N and C terminal interactors shows 32 proteins interacting more with the N-terminus of Kv1.3 and 40 proteins interacting with the C-terminus of Kv1.3. **(E)** String analysis shows close associations with N-terminal interactors. Gene Ontology highlights that many of these proteins are associated with metabolic processing. **(F)** String Analysis and GSEA analysis show that many of the C-terminal interactors are associated with signal transduction. Darker colors represent statistical significance in GSEA results, lighter color indicates an interactor that is not present in GSEA lists but interacting with Kv1.3, and edge thickness represents confidence of interactions based on literature. Differential Abundant proteins were calculated using paired t-test, where log P-value > 1.3 and Log_2_ Fold Change (FC) of +/-1 were considered significant. n=3 **p < 0.01, *** p< 0.001

We then performed differential enrichment analysis (DEA) to identify proteins that interact with Kv1.3 that appear both in the N and C terminal proteomes (*i.e.* proteins within labeling radius of TurboID) as well as proteins that show selective interactions with the N or C terminus of Kv1.3. Differential enrichment showed 1,681 N-terminal interacting proteins (**Fig. 2B**) and 1716 C-terminal interacting proteins (**Fig. 2C**). A large proportion of these N-term and C-term interactomes overlapped (**Supp.** Fig. 1), probably indicative of general Kv1.3 channel protein interactors that were within the labeling radius of TurboID, regardless of N-term or C-term domain specificity. While the N and C terminus of one Kv1.3 monomer may be separated in space, N and C termini of the adjacent Kv1.3 mononers in the tetrameric complex are likely closer to each other potentially explaining this overlap. The surprisingly large numbers of Kv1.3 interactors is also likely due to TurboID biotinylating proteins at all steps of Kv1.3 being processed, through the ER before being localized to the cell membrane. Accordingly, many of the interactors listed were associated with ER processing of proteins (*e.g.* SEC24A, SEC16A, HSPA5).

Although the majority of the N-terminal and C-terminal interactomes of Kv1.3 overlapped, some proteins were uniquely enriched in a domain-specific manner. DEA comparing N-terminal and C-terminal interactors directly, showed 32 proteins associated with the N terminus of Kv1.3 and 40 proteins associated with C-terminus of Kv1.3 (**Fig. 2D**). Gene set enrichment analysis (GSEA) highlighted biological processes associated with the N-terminus, involved in metabolic processing (*e.g.* TNXL1, ACSL1, and EPHX2) (**Fig. 2D**). The proteins that interacted with the N-terminus were likely associated with processing proteins to the ER such as some of the key metabolic interactors. In contrast, the C-term-specific interactome of Kv1.3 was enriched in proteins associated with signal transduction (*e.g.* SMAD3, RAC1, and CNOT1) (**Fig. 2D**). We also used Search Tool for the Retrieval of Interacting Genes/Proteins (STRING), to visualize known protein-protein interactions (PPIs) within these domain-specific Kv1.3 interactors (**Fig. 2E and F**). STRING analysis showed that Kv1.3 N-terminal interactors were involved in processing of proteins from the endoplasmic reticulum to the cell membrane *e.g.*. SRRM2, NACA, and TXNL1(**Fig. 2E**). Kv1.3 C-terminal interactors were largely associated with cell signaling (**Fig. 2F**). These results in HEK-293 cells indicate that distinct protein interactors of Kv1.3 channels can be defined using N and C terminal TurboID fusion, without significant disruption of channel physiology. Our analyses of both the protein lists and GSEA suggest that the N-terminus of Kv1.3 is primarily associated with protein processing whereas the C-terminus is associating with signaling proteins. Based on these studies in HEK-293 cells, we next applied TurboID-based proximity labeling to examine Kv1.3 channel interactomes in microglia.

### Generation and validation of stably transduced BV-2 microglial lines expressing N and C term Kv1.3-TurboID fusions

We chose the mouse-derived BV-2 microglial cell line as a model system to first identify proteins that interact with Kv1.3 channels in a N and C term-specific manner, and then test whether these domain-specific interactions were altered when microglia adopt pro-inflammatory phenotypes. We selected BV-2 cells, an immortalized murine microglia cell line for reasons detailed in the section Limitations of the Study below. BV-2 cells were transduced for 24h with lentiviruses (LV) or no LV for control, encoding one of three constructs confirmed in HEK-293 cells: a Kv1.3 Fusion protein with TurboID on the N-terminus, Kv1.3 fusion with TurboID on the C-terminus, and an Kv1.3-TurboID C-terminal fusion with the PDZ-binding domain removed (**Fig. 3A**). TurboID globally expressed in the cytoplasm was used as a positive control and untransduced BV-2 cells were utilized as a negative control. Following puromycin selection for seven days, transduced cell lines were either exposed to lipopolysaccharide (LPS, 100ng/mL) or PBS for 24h and then exposed to biotin for 1h to increase the stringency of the interactomes of Kv1.3 (**Fig. 3A**). LPS induction was used to induce a pro-inflammatory like state in BV-2 microglia. We first performed qRT-PCR and confirmed that Kcna3 mRNA was highly and comparably transcribed across all three lines as compared to sham transduced/control BV-2 cells (**Fig. 3B**). Western blot and flow cytometry showed biotinylation in Kv1.3-TurboID transduced microglia, which was absent in control BV-2 cells (**Fig. 3C and D**).

**Figure 3:**
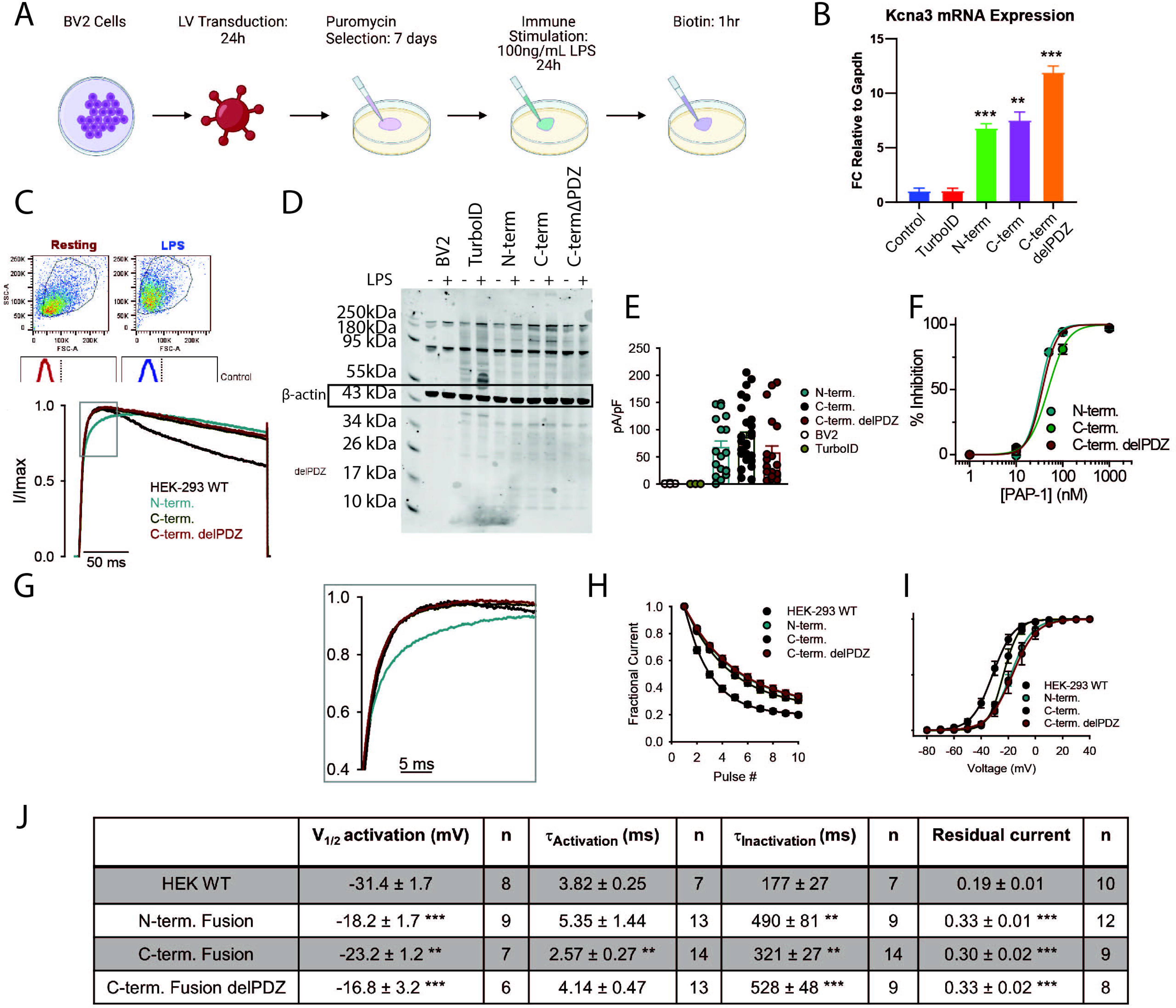
BV-2 cells transduced with Kv1.3-TurboID constructs show presence of biotinylation and channel activity. **(A)** Schematic of experimental design. BV-2 cells were transduced with Kv1-3 constructs described in Fig 1B and selected for plasmid uptake using puromycin. Cells were then exposed to LPS to induce an inflammatory response and biotin to allow for biotinylation of proximal proteins. **(B)**qRT-PCR of *Kcna3* transcript shows increased *Kcna3* mRNA expression in Kv1.3-TurboID transduced cell lines compared to control and *Gapdh*. Significance was calculated utilizing paired t-test. * means P < 0.05, **means P < 0.01, *** means P < 0.001. **(C)** Flow cytometry of biotinylated proteins using Streptavidin-488 shows high biotinylation in Kv1.3-TurboID transduced cell lines independent of LPS exposure. **(D)** Western blot depicting streptavidin-680 labeling, which highlights biotinylation increase in the presence of TurboID. Ponceau staining shows no change in protein concentration across samples. **(E)** Electrophysiology shows increased channel presence with transduction of Kv1.3-TurboID constructs, confirming successful transduction of the functional Kv1.3 channel. **(F)** Electrophysiology highlights inhibition utilizing PAP1, a Kv1.3 inhibitor, with transduction of Kv1.3-TurboID constructs. **(G)** Inactivation curves of electrophysiology data show a minor reduction of inactivation with Kv1.3-TurboID cells. **(H)** Fractional currents of Kv1.3 are slightly increased in Kv1.3-TurboID transduced cells. **(I)**Voltage inactivation of Kv1.3 is unchanged by transduction of Kv1.3-TurboID constructs. **(J)** Table summarizing electrophysiology results.

To determine whether stably transduced BV-2 lines exhibited functional Kv1.3 channel currents on the cell surface, we performed electrophysiological studies confirming that comparable and high Kv1.3 current densities were present across all cell lines (**Fig. 3E**). Similar to results observed in HEK-293 cells, BV-2 cells transduced with either of the three constructs fused with TurboID exhibited a positive shift in the voltage-dependence of activation, less use-dependent current decay, and delayed inactivation kinetics (**Fig. 3G-J**). The C-terminal fused Kv1.3 also showed enhanced activation kinetics compared to positive controls of HEK-293 cells transfected with unmodified Kv1.3 (**Fig. 3G and J**). PAP-1 also blocked Kv1.3 currents [37] (**Fig. 3I and Sup.** Fig. 2A) confirming the identity of the overexpressed Kv1.3 channels in BV-2 cells. These experiments showed that Kv1.3 is functionally active at comparable levels across all BV-2 Kv1.3-TurboID cell lines, with small yet significant increase in current with BV-2 cells transduced with Kv1.3 compred to a HEK_293 cell. These studies therefore lay the foundation for proteomic assessments of Kv1.3 channel interactomes in microglia using the proximity-labeling approach.

### Identification of Kv1.3 channel domain-specific protein interactors and molecular pathways in BV-2 microglia

In order to evaluate proteins that interact with Kv1.3 in microglia, biotinylated proteins were enriched from whole cell lysates of BV-2 lines, by affinity purification with streptavidin magnetic beads. Affinity purified proteins were verified by Western Blot and silver stain (**Supp.** Fig 2B). Via PCA of the cell lysates, we confirmed that LPS indication accounts for 23% of the variance of proteins in the total samples, whereas the addition of Kv1.3 does not alter variance within the samples (**Supp.** Fig. 3A). This indicated, overexpression of Kv1.3 did not significantly alter proteins present in each sample. DEA of proteins present in the cell lysates highlighted the LPS induced an increase of proteins associated with a neuroinflammatory response, including TLR2, IL-1α, and STAT1 (**Supp.** Fig. 3B). TurboID-normalized AP samples (n=3) were clustered using Principal component analysis (PCA). PC1 accounted for 65% variance and separates the samples with TurboID present from the un-transduced BV-2 controls as well as separated N-terminal fusions from C-terminal fusions (**Fig. 4A**). Separation of controls and all TurboID samples indicated that both biotinylation and enrichment was successful. PC2, accounted for 10% of the variance and separated the different cell types: overexpression of Kv1.3, un-transduced control, and global TurboID, which acts as a positive control for biotinylation but does not have Kv1.3 attached (**Fig. 4A**). This highlighted that the proteins biotinylated in BV-2 cells transduced with Kv1.3 are unique compared to un-transduced controls or the biotinylated proteome of BV-2 cells with global (cytosolic) TurboID localization.

**Figure 4:**
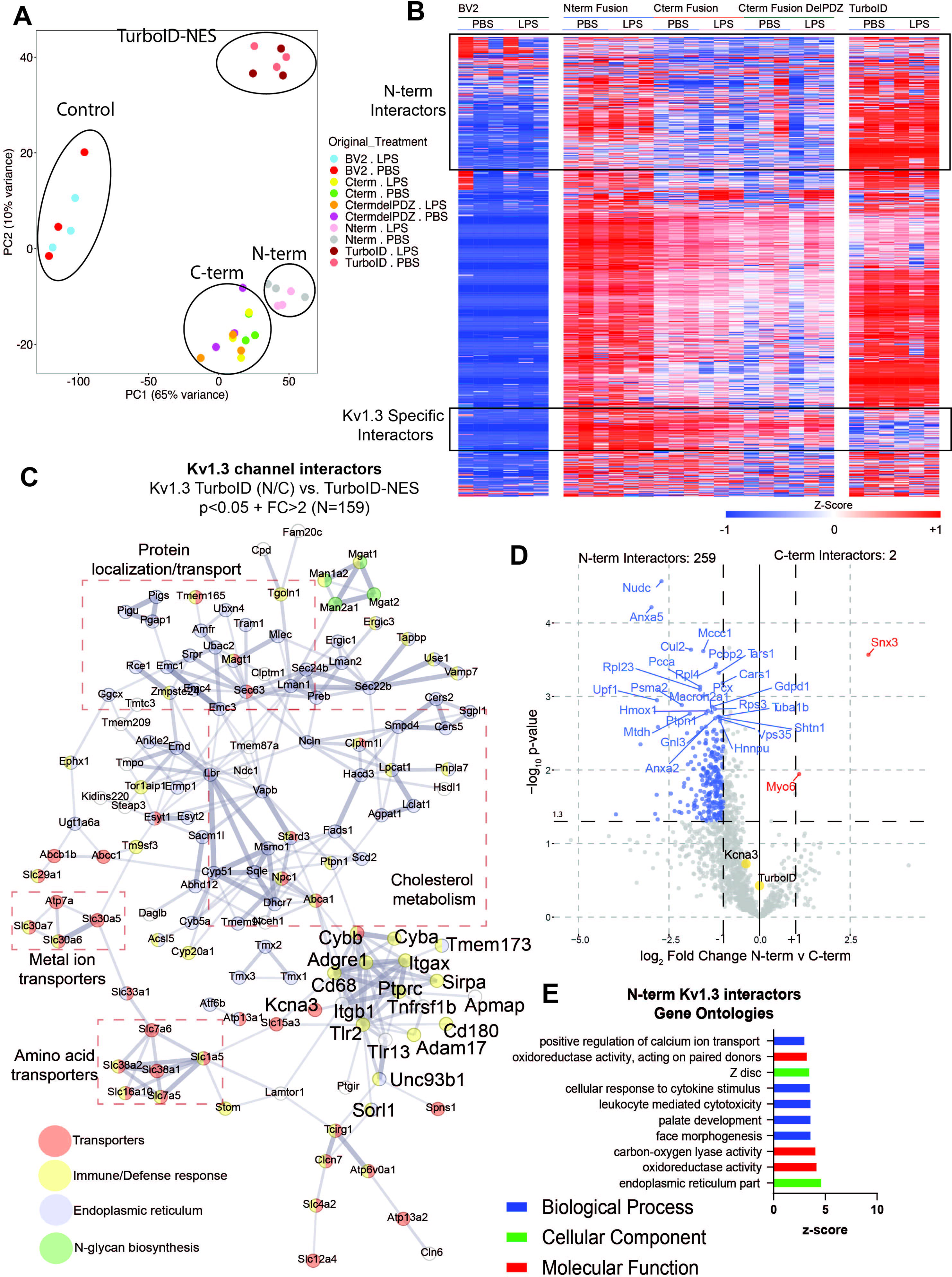
Microglial Kv1.3 have distinct N-terminal and C-terminal interactors. **(A)** Principal Component Analysis (PCA) of mass spectrometry of biotinylated proteins show distinct clustering of controls, TurboID, Kv1.3 N-term fusion, and Kv1.3 C-term fusion. **(B)** Heat map of biotinylated proteins, designed by Morpheus, show distinct clusters of Kv1.3 specific interactors, Kv1.3 N-terminal specific interactors, and Kv1.3 C-terminal interactors. Individual proteins were colored based on z-score, where the darker shades of red indicate +1 and the darker shades of blue indicates -1. Hierarchical clustering arranged proteins based on groups. **(C)** STRING analysis highlights proteins differentially interacting with Kv1.3 over global TurboID expression. Color indicates protein type. Associated protein groups are boxed. **(D)** Differential Enrichment Analysis (DEA) of proteins enriched with Kv1.3 N-terminal fusion and Kv1.3 C-terminal fusion indicates about 250 N-terminal interactors and 2 C-terminal interactors. **(E)** Gene set enrichment analysis (GSEA) analysis shows most of the interacting proteins with the N-terminus of Kv1.3 appear to be a part of calcium transport and oxidoreductase activity. Differential Abundant proteins were calculated using paired t-test, where log P-value > 1.3 and Log_2_ Fold Change (FC) of +/-1 were considered significant. n=3

We hypothesized that Kv1.3 would have distinct interactors between the N and C termini of the channel in microglia. There were many proteins present in the samples transduced with Kv1.3-TurboID that were absent in the untransduced controls (**Fig. 4B**). In total 991 proteins interacted either with the N or C terminus of Kv1.3 and were distinctly present in Kv1.3-TurboID transduced (**Fig. 4B and Supp.** Fig. 3 **C-E**). GSEA and STRING analyses of these Kv1.3-specific interactors identified protein localization/transport, metal ion transporters, cholesterol metabolism, amino acid transporters, N-glycan biosynthesis and immune/defense proteins (**Fig. 4C**). The most represented pathway in the Kv1.3 interactome in KEGG 2021 was protein processing in endoplasmic reticulum (corrected p 4.84E-22). These include ER translocon and translocation machinery, such as SEC61A1, SEC61B, SEC61G, SEC63, HSPA5. The largest interaction groups with Kv1.3 in general were associated with cholesterol metabolism (*i.e.* NPC1 and ABCA1) and protein localization/transport (*i.e.* Tmem165 and MAGT1) (**Fig. 4C**). We also identified a immune/defense response cluster of Kv1.3 channel interactors including integrins (ITGB1, ITGAX), TLRs (TLR2, TLR13), receptor tyrosine phosphatases (eg. PTPRC/CD45), lysosomal protein CD68 and Alzheimer’s disease related proteins (eg. SORL1, ITGAX) (**Fig. 4C**). Next, we compared how Kv1.3 terminal interactors differ between the termini.

Comparing the Kv1.3 N-terminus to C-terminus in BV-2 transduced cells, identified 284 proteins that preferentially interacted with the Kv1.3 N-terminus. These included proteins like TIMM50, a mitochondrial translocase protein, and NUDC and TXNL1, associated with protein processing (**Fig. 4D**). In comparison, far fewer exclusive C-term interactors were identified, including SNX3 and MYO6, both associated with intracellular processing (**Fig. 4D**). GSEA of the Kv1.3 N-terminal interactors show processes associated with calcium ion transport and oxidoreductase activity (**Fig. 4E**). In contrast to N and C terminal interactors in HEK-293 cells (albeit human), the N terminal interactome of Kv1.3 in BV-2 cells was much larger while the C terminal interactome was smaller. Our analyses in BV-2 cells identified the distinct Kv1.3 N-terminal and C-terminal interactors, among which the C terminal interactome was most impacted by LPS treatment of BV-2 cells, indicative of context-dependent changes of the Kv1.3 channel interactome.

### Pro-inflammatory activation preferentially modifies the C-term interactome of Kv1.3 channels in microglia

Microglia are well known to adopt distinct inflammatory profiles when exposed to immune stimuli, and their cellular functions are context-dependent [38, 39]. We therefore next assessed whether LPS pro-inflammatory activation impacted the Kv1.3 channel interactome in a domain-specific manner. We found that many C-terminal interactors were only identified as interactors when LPS was present that additionally present during an LPS response in cell lysates, although the N-terminal interactome was not impacted by LPS treatment (**Fig. 4B**). In the presence of LPS, we hypothesized the interactors of Kv1.3 would shift toward an inflammatory signaling. DEA showed negligible effects of LPS on the Kv1.3 N-terminal interactome, even though LPS effects were noted on the whole cell proteome (**Fig. 5A**). DEA showed in the presence of LPS there were 36 proteins increased and 27 proteins decreased in the C-terminal interactome (**Fig. 5B**). Proteins with increased interaction with the C terminus upon LPS activation included STAT1, C3, and TLR2, often associated with a proinflammatory response in microglia, whereas the decreased proteins include SNX3 and HSPA9. GSEA showed the C-terminal proteins that interacted with Kv1.3 in the absence of LPS, associated with the ER and transport (**Fig. 5C**). In the presence of LPS, Kv1.3 C-terminal interactors transitioned towards inflammatory proteins and immune effector processes (**Fig. 5D**). This increase in C terminal interactors by LPS cannot be explained by increased total protein abundance itself, because this increase was largely absent in the N-terminus as well as in the global TurboID conditions. In addition, what makes this interactome even more interesting is that there is a paucity of secreted proteins in the unstimulated Kv1. 3 interactome which suggested that these results may be selective. In fact, the overlap between the Kv1.3 interactome and the human secretome is just 63 proteins. Therefore, we conclude that the pro-inflammatory context of microglial activation specifically modifies the C-terminal interactome of Kv1.3 channels, potentially linking channel function with immune signaling machinery.

**Figure 5:**
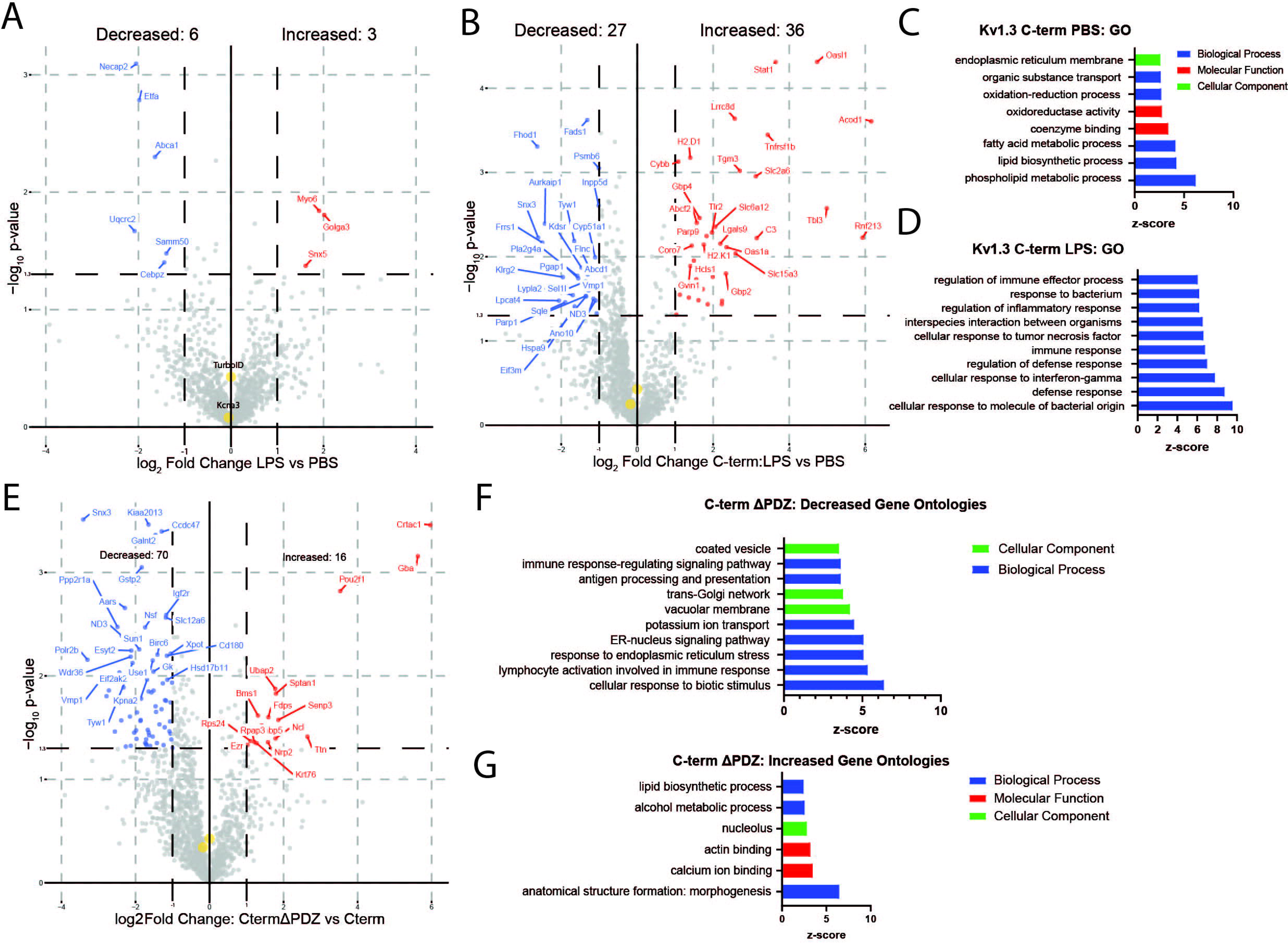
Inflammatory exposure results in a PDZ-binding domain dependent interactions with the C-terminus of Kv1.3. (A) Differential Enrichment Analysis (DEA) between N-terminal interactors and N-terminal interactors with LPS exposure highlights minimal change in N-terminal interactors with LPS exposure. (B) DEA between C-terminal interactors and C-terminal interactors show 27 proteins interacting with the C-term of Kv1.3 during homeostasis and 36 proteins interacting with the C-terminus during an LPS-induced inflammatory response. (C) Gene Set Enrichment Analysis (GSEA) analysis shows protein transport and processing terms downregulated during an LPS response. (D) GSEA shows an upregulation of immune signaling terms associated with the C-terminus of Kv1.3 during LPS immune stimulation. (E) DEA comparison of Kv1.3 C-terminal interactors and C-terminal interactors with the PDZ-binding domain removed shows 70 proteins downregulated and 16 upregulated with deletion of the PDZ-binding domain. (F) GSEA analysis of proteins downregulated with the deletion of the PDZ-binding domain are mostly associated with immune-signaling response and protein packaging. (G) GSEA analysis of proteins upregulated with deletion of the Kv1.3 C-terminal PDZ-binding domain shows terms associated with lipid biosynthesis and alcohol metabolism. Differential Abundant proteins were calculated using paired t-test, where log P-value > 1.3 and Log_2_ Fold Change (FC) of +/-1 were considered significant.

### Kv1.3 C-terminal inflammatory interactors dependent on the PDZ binding domain

The Kv1.3 PDZ-binding domain has been shown to be essential to Kv1.3 function [23]. Since the C-term of Kv1.3 interacted with signaling proteins across two mammalian cell lines, we hypothesized that deletion of the PDZ binding domain of the C-terminus of Kv1.3 would result in a reduction of interacting proteins with the C-terminus. DEA identified 70 Kv1.3 C-terminus interacting proteins with reduced interactions with the C-terminus upon deletion of the PDZ-binding domain, including SNX3, ND3, and NSF. Conversely, we found 16 proteins with increased interactions with the C-terminus when the PDZ-binding domain was removed, including GBA, NRP2, and CRTAC1 (**Fig. 5E**). GSEA of the proteins that interacted with the C-term in a PDZ-dependent manner (proteins with reduced interactions with Kv1.3 upon PDZ-binding domain removal), showed an association with the cellular component of coated vesicles and the biological processes of antigen processing and presentation and immune-response-regulating signaling pathway (**Fig. 5F**). Proteins that showed increased C-terminal interaction upon PDZ-binding domain removal were enriched in ontologies related to lipid biosynthesis and alcohol metabolic processing (**Fig. 5G**). These results highlighted several proteins and pathways related to the C-terminal Kv1.3 channel interactome that require the PDZ binding domain to interact with Kv1.3.

### Kv1.3 channel is present in mitochondria-enriched fractions in microglia

It has been reported that Kv1.3 channel can be detected at the plasma as well as the inner mitochondrial membranes in lymphocytes and cancer cells [15, 40, 41]. To assess the presence of Kv1.3 in mitochondria from BV-2 cells transduced with Kv1.3-TurboID constructs, we prepared subcellular fractions from total cell homogenates (Fraction 0) by sequential centrifugation resulted in Fraction A1 contained light membranes, Fraction A2 contained crude mitochondria, and Fraction A3 enriched in mitochondria (**Fig. 6A**). Western blot analysis of the mitochondrial marker HSP60 showed that HSP60 levels were increased in Fractions A3 compared to Fractions 0, confirming mitochondrial enrichment of fraction A3 was successful (**Fig. 6B**). Next, western blot analysis of these fractions, probed for V5 (fused to Kv1.3-TurboID or to TurboID), detected V5 bands that corresponded to predicted molecular weights of fusion proteins in all subcellular fractions from BV-2 cells transduced with Kv1.3-TurboID constructs (N-term, C-term, and C-term ΔPDZ), including the mitochondria-enriched fractions A3 (**Fig. 6C**). Small variabilities between V5 tagged N-terminus and C-terminus intensities may be due to small variability in TurboID expression. No bands were detected in un-transduced BV-2 cells (Neg. Control) (**Fig. 6C**). STRING analysis of Kv1.3 interactors that a present in the MITOCARTA 3.0 database data show 73 proteins associated with the mitochondria (**Fig. 6D**). These include proteins associated with transport of proteins to the mitochondria and mitochondrial ribosomal machinery (*e.g.* TIMM50, MRPS30, and TMX1). Our results provide evidence for mitochondrial Kv1.3 protein presence in mitochondrial fractions from BV-2 cells transduced with Kv1.3-TurboID constructs.

**Figure 6.**
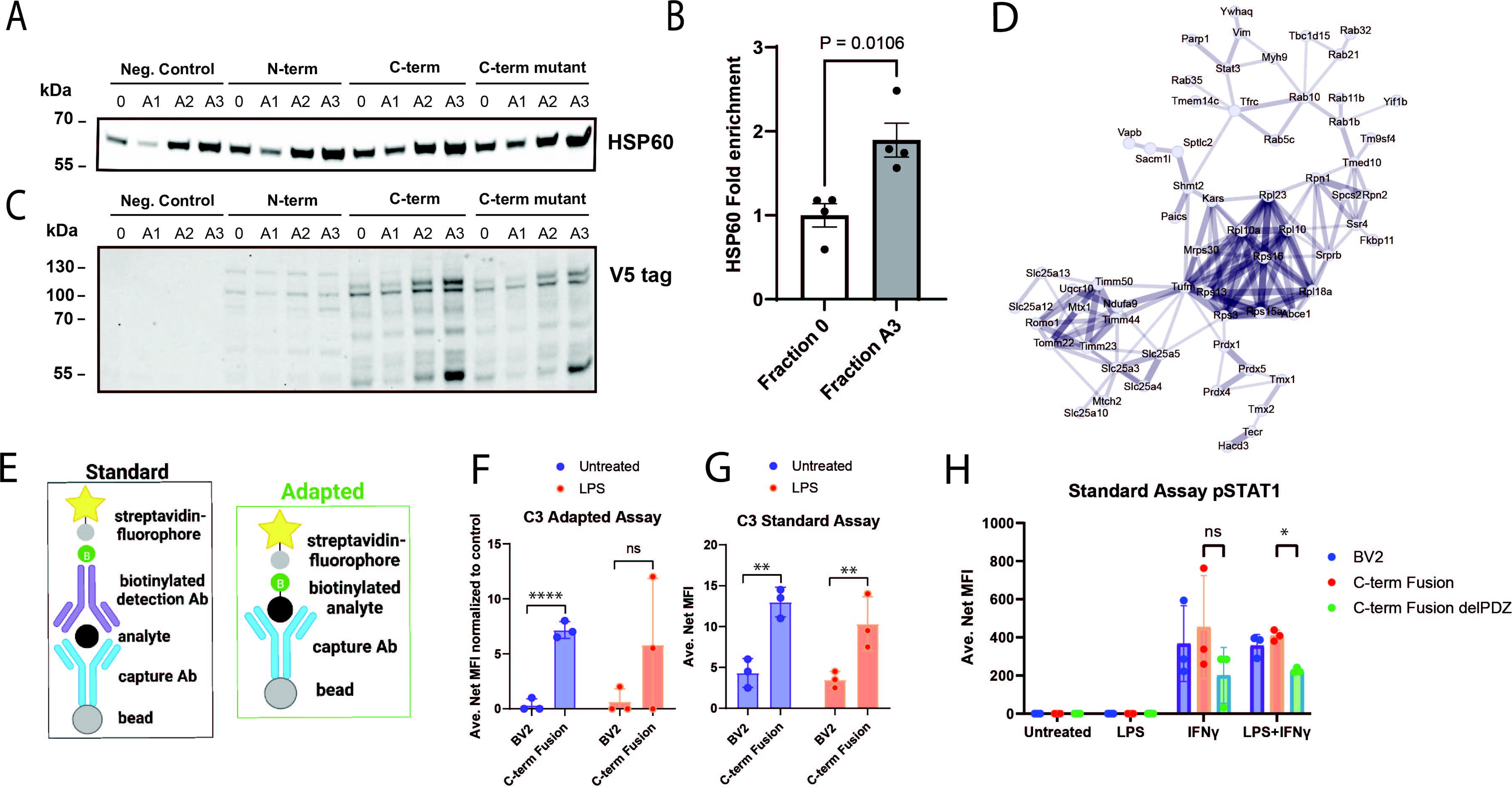
Confirmation of Kv1.3 interactions with mitochondria and immune signaling. **(A)** Validation of mitochondria-enrichment in the subcellular fractions obtained during the isolation process from untransduced BV-2 cells (Neg. control) and BV-2 cells transduced with Kv1.3-TurboID constructs (N-term, C-term, and C-term Mutant). Total homogenates (0) were fractionated to heavy membranes (A1), crude mitochondria (A2), and mitochondria-enriched fractions (A3) by sequential centrifugation; and then analyzed by Western blot using HSP60 as a mitochondrial marker. **(B)** Quantification of HSP60 comparing total homogenates (0) to mitochondria-enriched fractions (A3). P-value for two-sided unpaired T-test is indicated. **(C)** Representative blot of V5 tag (TurboID fusion) in fractions 0 – A3 from BV-2 cells transduced with Kv1.3-TurboID constructs. **(D)** STRING analysis of Kv1.3 interactors cross referenced to MITOCARTA 3.0 database highlights many proteins interacting with Kv1.3 associated with the mitochondria and the functional and physical interactions. **(E)** Schematic of standard and adapted Luminex assay. Standard assay includes a bead attached to a capture antibody that binds to the protein of interest, then a biotinylated antibody binds to the protein of interest to form a sandwich. Streptavidin with a fluorophore binds to the biotin. This estimates how much of a protein is in a sample. The adapted assay includes a capture antibody with bead and then adds streptavidin with the fluorophore directly to the protein of interest. The adapted assay captures abundance of proteins interacting with Kv1.3 directly. **(F)** Adapted Luminex assay of intracellular C3 shows an increased interaction with Kv1.3 at a resting state. **(G)** Standard Luminex assay of proteins isolated from BV-2 cells transduced with Kv1.3-TurboID highlights an increase in intracellular C3 with the overexpression of Kv1.3, independent of LPS exposure. **(H)**Standard Luminex assay of pSTAT1 shows activation of pSTAT1 in the presence of INF-γ, a known activator of the STAT1 signaling pathway. In the presence of LPS and INF-γ, the removal of the PDZ-binding domain in Kv1.3 leads to a decrease in the presence of pSTAT1. Significance was calculated utilizing paired t-test. * p < 0.05, **p < 0.01, ***p < 0.001, ****p < 0.0001. n=3

### Verification of interactions of Kv1.3 with C3 and pSTAT1 via the C terminal domain

Among the immune-related protein interactors of Kv1.3 that were measured by our MS studies, we nominated C3 and STAT1 as proteins of interest based on their known role in several neuroinflammatory and neurodegenerative diseases [42–47]. We used Luminex to validate these two Kv1.3 C terminal interactors, relevant to microglial immune functions, namely C3 and pSTAT1. The standard Luminex approach immobilizes the protein of interest on to a bead via capture antibody, and then uses a biotinylated detection antibody followed by streptavidin-fluorophore conjugate to quantify abundance of the target protein [31]. We adapted this approach by omission of the detection antibody, so that all C3/pSTAT1 would be captured but only their biotinylated forms would be detected, which proved a direct quantification of the biotinylated forms of these proteins from a total cell lysate without enrichment (**Fig. 6E**).

The adapted assay was able to detect biotinylated C3 in cell lysates from the C-term TurboID Kv1.3 BV-2 cells, regardless of LPS stimulation, confirming the MS result of an interaction between the C-term of Kv1.3 and C3 (**Fig. 6F**). In comparison to the adapted assay which only detects biotinylated C3 in cell lysates, the standard assay for C3 also showed an overall increased level of C3 in C-terminus Kv1.3 independent of global expression of TurboID(**Fig. 6G**). This finding can be explained by either increased C3 protein abundance in microglia when Kv1.3 is over-expressed, or, an increased C3 signal due to detection of the protein via the capture antibody as well as detection of the biotinylated C3 via streptavidin-fluorophore. Importantly, the adapted C3 levels detected in cell lysates from C-term TurboID Kv1.3 BV-2 cells, were greater than 50% of the C3 signal from the standard assay. This suggested that majority of C3 in BV-2 TurboID cells is biotinylated by TurboID, consistent with C3 being identified as an important Kv1.3 interactor. Since C3 is not traditionally a membrane-attached protein, this result most likely represents an interaction between Kv1.3 and C3 in the processing stage of C3 before its secretion, potentially at the stage of processing in the ER translocon. This was consistent with several ER translocon proteins identified as Kv1.3 interactors (eg. SEC61A1, SEC61B, SEC61G, SEC63, HSPA5), and enrichment of protein processing in the ER, as a major over-represented pathway in the Kv1.3 interactome.

We also measured levels of phosphorylated Stat1 (pStat1 Tyr 701) to reflect activated Stat1 in BV-2 cells. Stat1 phosphorylation is triggered by type 1 and 2 interferons, leading to activation of interferon-related gene expression which are important in anti-viral immune responses, as well as in neurological diseases [48]. Previously, a functional relationship between Kv1.3 channel activity and pStat1 phosphorylation was also suggested [9]. Our MS studies suggested that the C-term of Kv1.3 interacts with STAT1using the Luminex assay to detect pStat1 levels in cell lysates from BV2 cells, using LPS as a pro-inflammatory condining stimulus, and IFNγ as an activator of Type II interferon signaling. We also examined whether the PDZ-binding domain is required for this interaction. Standard Luminex measurements of pStat1 showed that IFNγ-treated BV-2 cells, regardless of pre-incubation with LPS, responded via increased pStat1 levels to IFNγ, and this effect of IFNγ was decreased by half when the PDZ-binding domain of the C-terminus Kv1.3 was deleted (**Fig. 6H**). This result supported an important role of the C-terminal domain, particularly the PDZ-binding domain, in the regulation of the interaction between Kv1.3 and STAT1 signaling proteins in microglia.

## DISCUSSION

### Kv1.3 interacts with immune signaling proteins in microglia, exhibiting domain-specific as well as context-dependent interaction patterns

Recent advances in the genetics of human neurological diseases and experimental manipulation of immune targets in disease models have strongly implicated microglia as causal mediators of disease pathogenesis across many neuroinflammatory and neurodegenerative diseases. Establishing microglial targets to neurological disease is essential to modern therapies. Kv1.3 appears to be a good candidate target due to it’s specificity to immune cells and dramatic shift in activity during an immune response. Kv1.3 physical interactions with immune proteins during a microglial activated state indicates that it is likely working in complex with other immune proteins. Further development of blockade strategies of Kv1.3 *in vivo* will help establish regulation of Kv1. 3 as a potential treatment. It is worthwhile to explore how inhibiting N and C terminal interactors may contribute to the formation of the initiation of immune signaling in microglia.

Among several immune targets for neuroimmune modulation in neurological diseases, the Kv1.3 channel has emerged as a promising target that is highly expressed in pro-inflammatory subsets of disease-associated microglia (DAM). To address gaps in our current understanding of how Kv1.3 channels regulate immune functions of microglia, we applied a proximity-labeling approach to label the protein-protein interactome of Kv1.3 channels in mammalian cells, including mouse microglia *in vitro*. The TurboID proximity labeling technique has emerged as a tool to determine what proteins are interacting with in a close proximity to other proteins. By fusing the biotin ligase TurboID to the C or N terminus of Kv1.3 channels in microglia, we labeled the C and N terminal interactomes of Kv1.3 and used MS to quantify these biotinylated proteins after streptavidin-based enrichment. We identified over 900 proteins that interact with Kv1.3 channels in both HEK-293 cells and BV-2 microglia. Using a combination of electrophysiology, biochemical and immunofluorescence microscopy approaches, we confirmed that fusion of TurboID to the N or C terminal domains of Kv1.3 via a flexible linker, has minimal impact on channel localization and biophysical properties. During a homeostatic state in HEK-293 and BV-2 cells, it appears that the majority of interactors with Kv1.3 are indistinguishable between the N-terminus and the C-terminus but distinct from a global exposure to TurboID. LPS acts through TLR, which are involved in pathogenesis of both AD and PD [26, 27]. Using LPS allows a more precise, pathway-specific Kv1.3 interactome analysis. The Kv1.3 interactors are strongly associated with many disease-associated signaling pathways, including proteins like TLR2 and ITGAX, many of which are associated with immune responses and the DAM phenotype of microglia neurodegeneration [26, 27, 49]. Within MS-quantified Kv1.3 interacting proteins, we identified distinct groups of proteins that preferentially interact with N and C terminal domains of Kv1.3. When comparing the N-terminal interactors to the C-terminal interactors, it appears that the N-terminus is responsible primarily for interacting with proteins associated with protein processing while the C-terminus interacts with signaling proteins during an LPS response (**Fig. 7**).

**Figure 7.**
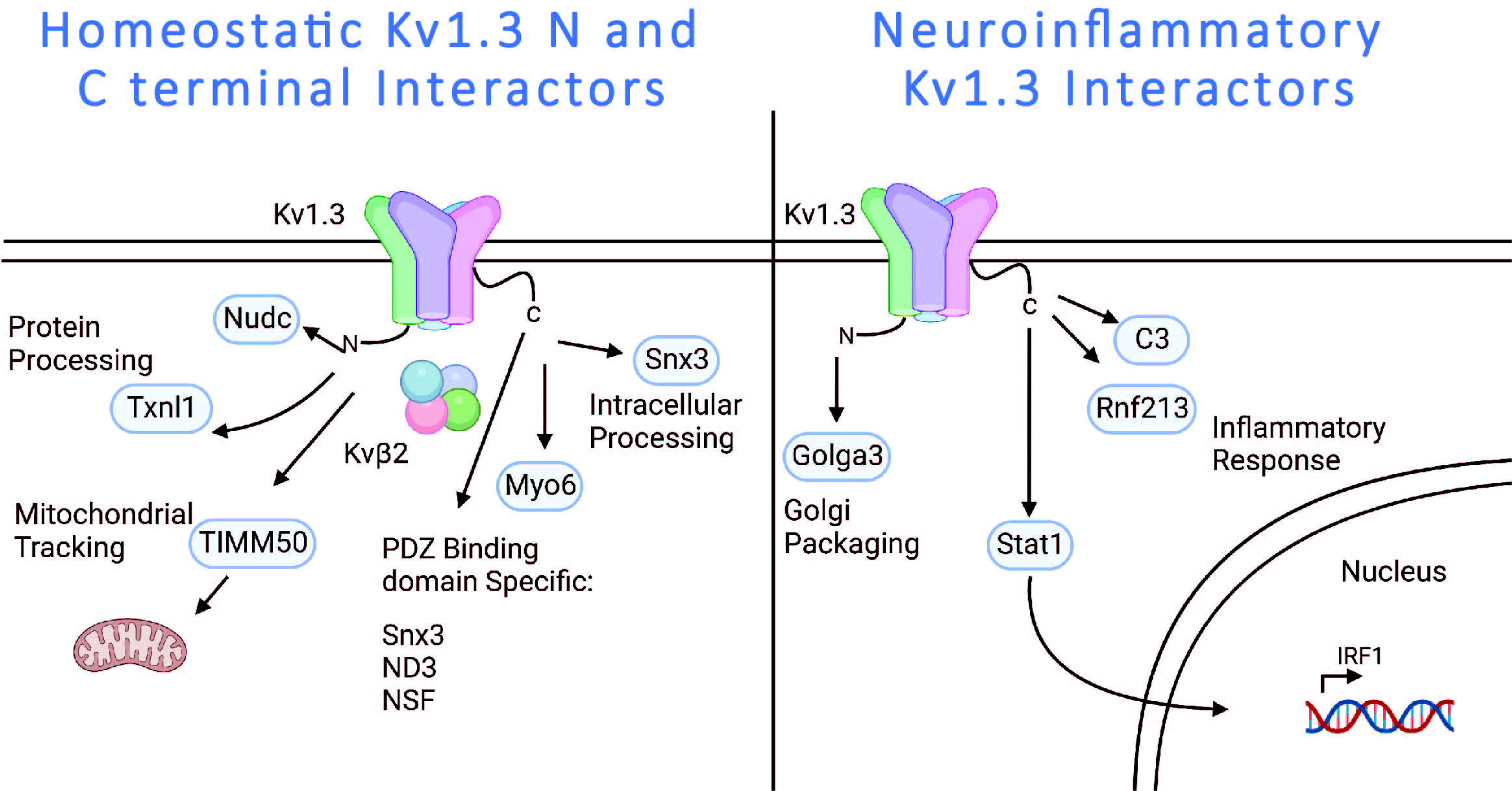
Schematic of Kv1.3 interactors in microglia during homeostatic and neuroinflammatory states. During homeostasis, many of the interactors of Kv1.3 are shared. The N-terminus appears to be responsible for protein processing and tracking to the mitochondria, whereas the C-terminus has some PDZ-binding domain specific interactors that are largely associated with intracellular processing. During LPS stimulation of an immune response, the N-terminus of Kv1.3 has very little change in the interacting partners, whereas the C-terminus has PDZ-binding domain dependent interactors associated with immune signaling.

Given the context-dependency of microglial states and functions in disease, we assessed whether the Kv1.3 channel interactome is altered when microglia adopt pro-inflammatory states, using LPS as a well-known pro-inflammatory stimulus [38, 39]. We found that while the N-terminal interactome was not impacted by LPS, the C-terminal interactome underwent significant reorganization, such that immune and signaling proteins showed higher levels of interaction, indicative of a domain-specific and context-specific effect of LPS. the proteins with higher interaction with the C terminus during LPS activation included immune signaling proteins and immune proteins that are part of the TLR and MHC complex, like STAT1 and C3 (**Fig. 7**). It is very likely that the presence and activation of the Kv1.3 increases pSTAT1 activity and aids in the induction of an inflammatory response in microglia. Similarly, in HEK-293 cells, the C-terminus of Kv1.3 also interacts with key signaling proteins such as MEK, a kinase that activates ERK signaling. We also found that a large fraction of the C terminal interactome is dependent on the PDZ binding domain [50]. Removal of the PDZ binding domain resulted in a reduction of immune signaling protein interactions, suggesting that this binding domain is necessary for physical interactions between Kv1.3 and these proteins (**Fig. 7**). Proteins such as STAT1 and C3 are independent of the PDZ-binding domain, whereas others like, CD190 and SUN1 are PDZ-binding domain dependent. While both status and the PDZ-binding domain interacting proteins are involved in signaling the PDZ-binding regulation does not influence the interaction of proteins during an activate state. This means that other domains of Kv1.3 on the C term (or another intermediary interactor) may be determining how immune status impacts C term interactions.

These traits of Kv1.3 provide strong insight on how Kv1.3 may regulate microglial immune functions, and how regulation of the Kv1.3 channel might be useful as a therapeutic target. Perhaps, the observed beneficial effects of blocking Kv1.3 channels in a plethora of neurological disease models may be explained in part by the interaction between Kv1.3 channels and immune signaling proteins that co-assemble in activated microglia.

### Kv1.3 presence and activity in the mitochondria

Our MS-based studies also suggested an interaction between Kv1.3 and mitochondrial proteins. Specifically, we found that the N terminal interactome of Kv1.3 includes proteins such as TIMM50, suggesting that some Kv1.3 channels in microglia are transported to the mitochondria, where they are likely present in the inner membrane[15, 51]. To validate this biochemically, we performed mitochondrial fractionation from whole cell lysates and found that Kv1.3 protein (detected via the V5 tag fused to Kv1.3) was indeed enriched in the mitochondrial fractions along with a known canonical mitochondrial marker (HSP60). Our finding of Kv1.3 protein presence in mitochondria of microglia aligns with prior observations that Kv1.3 channels can localize to the mitochondria in T cells, where they may regulate apoptosis, mitochondrial potential and proton flux. Future proteomics studies of the Kv1.3 channel interactome specifically in microglial mitochondria, and mechanistic studies investigating the functional implications of these interactors, are therefore warranted. These functional implications include, how Kv1.3 in the mitochondria is changed during LPS exposure and how Kv1.3 could regulate mitochondrial function during inflammation.

### Kv1.3 potassium channels may regulate immune function of microglia via protein-protein interactions

Microglia in neurological diseases have a higher expression of Kv1.3 [5, 7, 52]. Kv1.3 potassium channel activity is increased when membrane potential is reduced by the influx of calcium into microglia during an inflammatory response [53]. Activity of Kv1.3 clearly plays an important role in regulating immune function in microglia, however, it appears that Kv1.3 is also directly interacting with immune proteins independent of activity. This is evident by the fact that Kv1.3 has multiple binding locations including the SH3 binding region located on the N-terminus and the PDZ binding domain located on the C-terminus [30]. Previous studies have shown some immune signaling proteins directly interacting with Kv1.3, including MEK, a necessary part of the ERK activation pathway [50]. The ERK activation pathway is strongly linked to neurological diseases and is a necessary step in the activation of microglia. The interactions our paper highlights the role Kv1.3 plays in signaling beyond flow of potassium.

These findings strengthen previous connections between Kv1.3 and immune interactors specifically highlighting the potential for physical interactors of Kv1.3 with the TLR proteins in an immune stimulation complex. Prior work has also highlighted in mouse microglia cells (MMC), a murine cell line, as well as primary microglia which endogenously express Kv1.3 blockade of Kv1.3 using Shk-223 results in p38 and MAPK regulation[9, 54]. The current manuscript describes pStat1 is regulated by the C-terminus, previous data in our lab indicated that Stat1 phosphorylation is regulated by Kv1.3 function in BV-2, and Kv1.3 may interact with TLR[11]. This work highlights that the C terminal interactome has several immune proteins including TLR2, MHC proteins and STAT1, further supporting previous predictions[9]. These data and previous reports highlight that Kv1.3 channels may regulate or be functionally coupled with type 1/2 IFN signaling in microglia. IFN signaling is important in neuroinflammatory diseases like AD and aging, and in microglia, where IFN-based signatures have been identified in human brain[55, 56]. This provides rational for further evaluation of Kv1.3 influence in immune signaling *in vivo* and *in vitro*. Based on interactions with anchoring proteins (*e.g.* cortactin, integrins etc.), Kv1.3 likely becomes a part of a much larger macro-molecular complex of proteins that come together as an immune complex.

### Other Considerations

There are many desirable benefits to utilizing an *in vitro* system to gain insights into Kv1.3’s interactomes, including complete control over both the concentration and duration of stimulus in a highly pathway-specific manner to yield more precise results. However, quite a few limitations present. While BV-2 cells retained many properties to naïve to primary mouse microglia, they are not physiological replicate of *in vivo* microglia. In general, BV-2 cells are known to have adopted a more reactive phenotype compared to primary microglia, but do have similar inflammatory responses [46]. Endogenously, BV-2 cells do not express Kv1.3, however, our model utilized a transduced overexpression of Kv1.3. There is an inherent risk that the overexpression model leads to increase of proteins that could be interacting with Kv1.3 than in normal expression of Kv1.3. Similar issues arise when referring to the immortalized HEK-293 cell line, such as constant division and being derived from cancer cells. There is risk that the overexpression model leads to increase of proteins that could be interacting with Kv1.3 than in normal expression of Kv1.3.

TurboID is a proximity labeling technique, meaning that biotinylation of proteins can occur as long as it’s within the 10-30nm radius of labeling [24]. As a result, TurboID can biotinylate proteins that are just within proximity and not physically interacting with Kv1.3. Fortunately, the radius of labeling for TurboID fused with Kv1.3 should be within the range of direct and immediate interactors, including those are transient or fixed interactors. In addition, TurboID is can biotinylate proteins that are near Kv1.3 as it’s being packaged to the cell surface, such as labeling while Kv1.3 is present in the ER, which could explain the interactions observed with secreted proteins such as C3 [24]. In support of this, we found ER translocon/chaperone proteins such as HSP90 and HSPA5 as interactors of Kv1.3. It will also be important to determine whether these interactions are dependent or independent of K conductance via the pore of Kv1.3 channels. As in, does Kv1.3 initiate the formation of the immune signaling complex or is the activity of Kv1.3 a response to the formation of the immune signaling complex? With the establishment of the potential interactors of Kv1.3 in microglia, future studies are needed to determine how regulation of Kv1.3 alters in immune and disease response of microglia *in vivo* and how the directionality of immune interactions can be determined.

## CONCLUSIONS

In conclusion, we used proximity-based proteomics and identified several novel protein interactors of Kv1.3 channels in microglia, including domain specific interactors of the N terminus (*e.g*. TIMM50) and C terminus (*e.g.* STAT1 and C3) of the channel. While the N terminal interactome is larger and includes anchoring, localization and metabolic processing related proteins, the C terminal interactome in enriched in immune signaling proteins, and many of these are directly governed by the immune status of microglia, and some are dependent on the PDZ-binding domain.

## FIGURE LEGENDS

**Supplemental Figure 1: Confirmation of Hek-293 cell Kv1.3 function and TurboID activity. (A)** Traces of Kv1.3 electrophysiology of HEK-293 transfected with WT Kv1.3. **(B)** Traces of Kv1.3 activity of HEK-293 cells transfected with TurboID fused to the N-terminus of Kv1.3. **(C)** Traces of Kv1.3 activity of HEK-293 cells transfected with TurboID fused to the C-terminus of Kv1.3. **(D)** Ponceau staining of HEK-293 showing even protein loading across western blots prior to affinity purification. **(E)** Post-affinity purification western blot and silverstain show streptavidin labeling of proteins transfected with TurboID and proper affinity purification. **(F)** DEA comparison of HEK-293 Kv1.3 interactors on the N-terminus and C-terminus. Differential Abundant proteins were calculated using paired t-test, where log P-value > 1.3 and Log_2_ Fold Change (FC) of +/-1 were considered significant. n=3

**Supplemental Figure 2: Confirmation of BV-2 cell Kv1.3 function and TurboID activity. (A)** Traces of Kv1.3 electrophysiology of BV-2 cells transduced with TurboID fused to the N-terminus of Kv1.3. **(B)** Traces of Kv1.3 activity of BV-2 cells transduced with TurboID fused to the C-terminus of Kv1.3. **(C)** Traces of Kv1.3 activity of BV-2 cells transduced with TurboID fused to the C-terminus of Kv1.3 with the PDZ-binding domain removed. **(D)** Post-affinity purification western blot and silverstain show streptavidin labeling of proteins transfected with TurboID and proper affinity purification. n=3

**Supplemental Figure 3: BV-2 cells successfully produce inflammatory responses and cells transduced with Kv1.3 have distinct interactors. (A)** Principal Component Analysis (PCA) of mass spectrometry of BV-2 cell lysates shows distinct separation between cells exposed to LPS compared to PBS. There is little variance explained in the cell lysate proteomes that show difference between overexpression of Kv1.3. **(B)** DEA of whole cell lysates of BV-2 cells induced with LPS have increased presence of key inflammatory proteins. **(C)** DEA of affinity purified BV-2 cells with an N-terminal fusion of TurboID to Kv1.3 compared to controls shows presence of proteins interacting with the Kv1.3 channel. **(D)** DEA of affinity purified BV-2 cells with an C-terminal fusion of TurboID to Kv1.3 compared to controls shows presence of proteins interacting with the Kv1.3 channel. **(E)** DEA of affinity purified BV-2 cells with an C-terminal fusion of TurboID to Kv1.3 with a deletion of the PDZ-binding domain compared to controls shows presence of proteins interacting with the Kv1.3 channel. Differential Abundant proteins were calculated using paired t-test, where log P-value > 1.3 and Log_2_ Fold Change (FC) of +/-1 were considered significant. n=3

**Supplemental Figure 4: Full-length Western blot images of V5 tag in total homogenates (A) and mitochondria-enriched fractions (B).** Fractionation experiments were run in triplicates and TurboID was included as positive control for anti-V5 tag antibody.

## Supporting information

Supplemental Datasheet 1

Supplemental Datasheet 3

Supplemental Datasheet 2

Supplemental Datasheet 4

## Acknowledgements

Research from this publication was supported by the national institute of Aging of the National Institute of Health: 5T32GM135060-03 (CAB), 1F31AG074665-01 (CAB), R01NS114130 (SR), R01AG075820 (SR, NTS, LBW), R01 ES034796 (EW), and 1RF1AG060285 (VF). This research was supported by the Viral Vector Core of the Emory Center for Neurodegenerative Disease Core Facilities, the Emory Flow Cytometry Core (EFCC), the Custom Cloning Core Division of Emory Integrated Genomics Core (EIGC), and the Integrated Cellular Imaging Core (ICI) all at Emory University. Christina Ramelow for helping design a figure. Biorender for software to create images.

## REFERENCES

1. Woodburn, S.C., J.L. Bollinger, and E.S. Wohleb, The semantics of microglia activation: neuroinflammation, homeostasis, and stress. Journal of Neuroinflammation, 2021. 18(1): p. 258.

2. Schafer, D.P., et al., Microglia sculpt postnatal neural circuits in an activity and complement-dependent manner. Neuron, 2012. 74(4): p. 691–705.

3. Keren-Shaul, H., et al., A Unique Microglia Type Associated with Restricting Development of Alzheimer&#x2019;s Disease. Cell, 2017. 169(7): p. 1276–1290.e17.

4. Deczkowska, A., et al., Disease-Associated Microglia: A Universal Immune Sensor of Neurodegeneration. Cell, 2018. 173(5): p. 1073–1081.

5. Rangaraju, S., et al., Potassium Channel Kv1.3 Is Highly Expressed by Microglia in Human Alzheimer’s Disease. Journal of Alzheimer’s Disease, 2015. 44: p. 797–808.

6. Austin, S.A., et al., Alpha-synuclein expression modulates microglial activation phenotype. J Neurosci, 2006. 26(41): p. 10558–63.

7. Qin, C., et al., Dual Functions of Microglia in Ischemic Stroke. Neuroscience Bulletin, 2019. 35(5): p. 921–933.

8. Gray, S.C., K.J. Kinghorn, and N.S. Woodling, Shifting equilibriums in Alzheimer’s disease: the complex roles of microglia in neuroinflammation, neuronal survival and neurogenesis. Neural Regen Res, 2020. 15(7): p. 1208–1219.

9. Ramesha, S., et al., Unique molecular characteristics and microglial origin of Kv1.3 channel– positive brain myeloid cells in Alzheimer’s disease. Proceedings of the National Academy of Sciences, 2021. 118(11): p. e2013545118.

10. Sarkar, S., et al., Kv1.3 modulates neuroinflammation and neurodegeneration in Parkinson’s disease. The Journal of Clinical Investigation, 2020. 130(8): p. 4195–4212.

11. Gao, T., et al., Temporal profiling of Kv1.3 channel expression in brain mononuclear phagocytes following ischemic stroke. Journal of Neuroinflammation, 2019. 16(1): p. 116.

12. Selvakumar, P., et al., Structures of the T cell potassium channel Kv1.3 with immunoglobulin modulators. Nature Communications, 2022. 13(1): p. 3854.

13. Pérez-García, M.T., P. Cidad, and J.R. López-López, The secret life of ion channels: Kv1.3 potassium channels and proliferation. American Journal of Physiology-Cell Physiology, 2018. 314(1): p. C27–C42.

14. Wang, X., et al., Kv1.3 Channel as a Key Therapeutic Target for Neuroinflammatory Diseases: State of the Art and Beyond. Frontiers in Neuroscience, 2020. 13.

15. Capera, J., et al., The Mitochondrial Routing of the Kv1.3 Channel. Frontiers in oncology, 2022. 12: p. 865686–865686.

16. Di Lucente, J., et al., The voltage-gated potassium channel Kv1.3 is required for microglial pro-inflammatory activation in vivo. Glia, 2018. 66(9): p. 1881–1895.

17. Marks, D.R. and D.A. Fadool, Post-synaptic density perturbs insulin-induced Kv1.3 channel modulation via a clustering mechanism involving the SH3 domain. J Neurochem, 2007. 103(4): p. 1608–27.

18. Birkner, K., et al., β1-Integrin– and KV1.3 channel–dependent signaling stimulates glutamate release from Th17 cells. The Journal of Clinical Investigation, 2020. 130(2): p. 715–732.

19. Levite, M., et al., Extracellular K+ and Opening of Voltage-Gated Potassium Channels Activate T Cell Integrin Function: Physical and Functional Association between Kv1.3 Channels and β1 Integrins. Journal of Experimental Medicine, 2000. 191(7): p. 1167–1176.

20. Chen, X., et al., Structure of the full-length Shaker potassium channel Kv1.2 by normal-mode-based X-ray crystallographic refinement. Proc Natl Acad Sci U S A, 2010. 107(25): p. 11352–7.

21. Long, S.B., et al., Atomic structure of a voltage-dependent K+ channel in a lipid membrane-like environment. Nature, 2007. 450(7168): p. 376–382.

22. Tyagi, A., et al., Rearrangement of a unique Kv1.3 selectivity filter conformation upon binding of a drug. Proceedings of the National Academy of Sciences, 2022. 119(5): p. e2113536119.

23. Doczi, M.A., D.H. Damon, and A.D. Morielli, A C-terminal PDZ binding domain modulates the function and localization of Kv1.3 channels. Exp Cell Res, 2011. 317(16): p. 2333–41.

24. Cho, K.F., et al., Proximity labeling in mammalian cells with TurboID and split-TurboID. Nature Protocols, 2020. 15(12): p. 3971–3999.

25. Maes, M.E., et al., Targeting microglia with lentivirus and AAV: Recent advances and remaining challenges. Neurosci Lett, 2019. 707: p. 134310.

26. Venezia, S., et al., Toll-like receptor 4 deficiency facilitates α-synuclein propagation and neurodegeneration in a mouse model of prodromal Parkinson’s disease. Parkinsonism Relat Disord, 2021. 91: p. 59–65.

27. Zhou, Y., et al., TLR4 Targeting as a Promising Therapeutic Strategy for Alzheimer Disease Treatment. Front Neurosci, 2020. 14: p. 602508.

28. Sunna, S., et al., Cellular Proteomic Profiling Using Proximity Labeling by TurboID-NES in Microglial and Neuronal Cell Lines. Mol Cell Proteomics, 2023. 22(6): p. 100546.

29. Mahdavi, A., et al., Engineered Aminoacyl-tRNA Synthetase for Cell-Selective Analysis of Mammalian Protein Synthesis. J Am Chem Soc, 2016. 138(13): p. 4278–81.

30. Beeton, C., et al., A novel fluorescent toxin to detect and investigate Kv1.3 channel up-regulation in chronically activated T lymphocytes. J Biol Chem, 2003. 278(11): p. 9928–37.

31. Rayaprolu, S., et al., Cell type-specific biotin labeling in vivo resolves regional neuronal and astrocyte proteomic differences in mouse brain. Nat Commun, 2022. 13(1): p. 2927.

32. Olsson, A., et al., Single-cell analysis of mixed-lineage states leading to a binary cell fate choice. Nature, 2016. 537(7622): p. 698–702.

33. Emig, D., et al., AltAnalyze and DomainGraph: analyzing and visualizing exon expression data. Nucleic Acids Res, 2010. 38(Web Server issue): p. W755–62.

34. Szklarczyk, D., et al., The STRING database in 2023: protein-protein association networks and functional enrichment analyses for any sequenced genome of interest. Nucleic Acids Res, 2023. 51(D1): p. D638–d646.

35. Jiménez-Pérez, L., et al., Molecular Determinants of Kv1.3 Potassium Channels-induced Proliferation. J Biol Chem, 2016. 291(7): p. 3569–80.

36. Jürgen, K., Functional Expression Of GFP-Tagged Kv1.3 And Kv1.4 Channels In HEK 293 Cells. European Journal of Neuroscience, 1998. 10(12): p. 3908–3912.

37. Schmitz, A., et al., Design of PAP-1, a Selective Small Molecule Kv1.3 Blocker, for the Suppression of Effector Memory T Cells in Autoimmune Diseases. Molecular Pharmacology, 2005. 68(5): p. 1254–1270.

38. Butovsky, O., et al., Identification of a unique TGF-β-dependent molecular and functional signature in microglia. Nat Neurosci, 2014. 17(1): p. 131–43.

39. Belhocine, S., et al., Context-dependent transcriptional regulation of microglial proliferation. Glia, 2022. 70(3): p. 572–589.

40. Gulbins, E., et al., Role of Kv1.3 mitochondrial potassium channel in apoptotic signalling in lymphocytes. Biochim Biophys Acta, 2010. 1797(6-7): p. 1251–9.

41. Styles, F.L., et al., Kv1.3 voltage-gated potassium channels link cellular respiration to proliferation through a non-conducting mechanism. Cell Death Dis, 2021. 12(4): p. 372.

42. Qin, C., et al., Fingolimod Protects Against Ischemic White Matter Damage by Modulating Microglia Toward M2 Polarization via STAT3 Pathway. Stroke, 2017. 48(12): p. 3336–3346.

43. Rangaraju, S., et al., Identification and therapeutic modulation of a pro-inflammatory subset of disease-associated-microglia in Alzheimer’s disease. Molecular Neurodegeneration, 2018. 13(1): p. 24.

44. Qin, H., et al., Inhibition of the JAK/STAT Pathway Protects Against α-Synuclein-Induced Neuroinflammation and Dopaminergic Neurodegeneration. The Journal of Neuroscience, 2016. 36(18): p. 5144–5159.

45. Maier, M., et al., Complement C3 Deficiency Leads to Accelerated Amyloid β Plaque Deposition and Neurodegeneration and Modulation of the Microglia/Macrophage Phenotype in Amyloid Precursor Protein Transgenic Mice. The Journal of Neuroscience, 2008. 28(25): p. 6333–6341.

46. Butler, C.A., et al., Microglial phagocytosis of neurons in neurodegeneration, and its regulation. Journal of Neurochemistry, 2021. 158(3): p. 621–639.

47. Bourel, J., et al., Complement C3 mediates early hippocampal neurodegeneration and memory impairment in experimental multiple sclerosis. Neurobiology of Disease, 2021. 160: p. 105533.

48. Platanias, L.C., Mechanisms of type-I- and type-II-interferon-mediated signalling. Nature Reviews Immunology, 2005. 5(5): p. 375–386.

49. Muraoka, S., et al., Enrichment of Neurodegenerative Microglia Signature in Brain-Derived Extracellular Vesicles Isolated from Alzheimer’s Disease Mouse Models. J Proteome Res, 2021. 20(3): p. 1733–1743.

50. Kan, X.-H., et al., Kv1.3 potassium channel mediates macrophage migration in atherosclerosis by regulating ERK activity. Archives of Biochemistry and Biophysics, 2016. 591: p. 150–156.

51. Szabò, I., et al., A novel potassium channel in lymphocyte mitochondria. J Biol Chem, 2005. 280(13): p. 12790–8.

52. Stefanova, N., Microglia in Parkinson’s Disease. Journal of Parkinson’s Disease, 2022. 12: p. S105–S112.

53. Nguyen, H.M., et al., Biophysical basis for Kv1.3 regulation of membrane potential changes induced by P2X4-mediated calcium entry in microglia. Glia, 2020.

54. Fordyce, C.B., et al., Microglia Kv1.3 channels contribute to their ability to kill neurons. J Neurosci, 2005. 25(31): p. 7139–49.

55. Doorn, K.J., et al., Microglial phenotypes and toll-like receptor 2 in the substantia nigra and hippocampus of incidental Lewy body disease cases and Parkinson’s disease patients. Acta Neuropathol Commun, 2014. 2: p. 90.

56. Althafar, Z.M., Targeting Microglia in Alzheimer’s Disease: From Molecular Mechanisms to Potential Therapeutic Targets for Small Molecules. Molecules, 2022. 27(13).

