## Supplementary figures and images for "Proximity labeling proteomics reveals Kv1.3 potassium channel immune interactors in microglia"

### Supplemental Figure 1

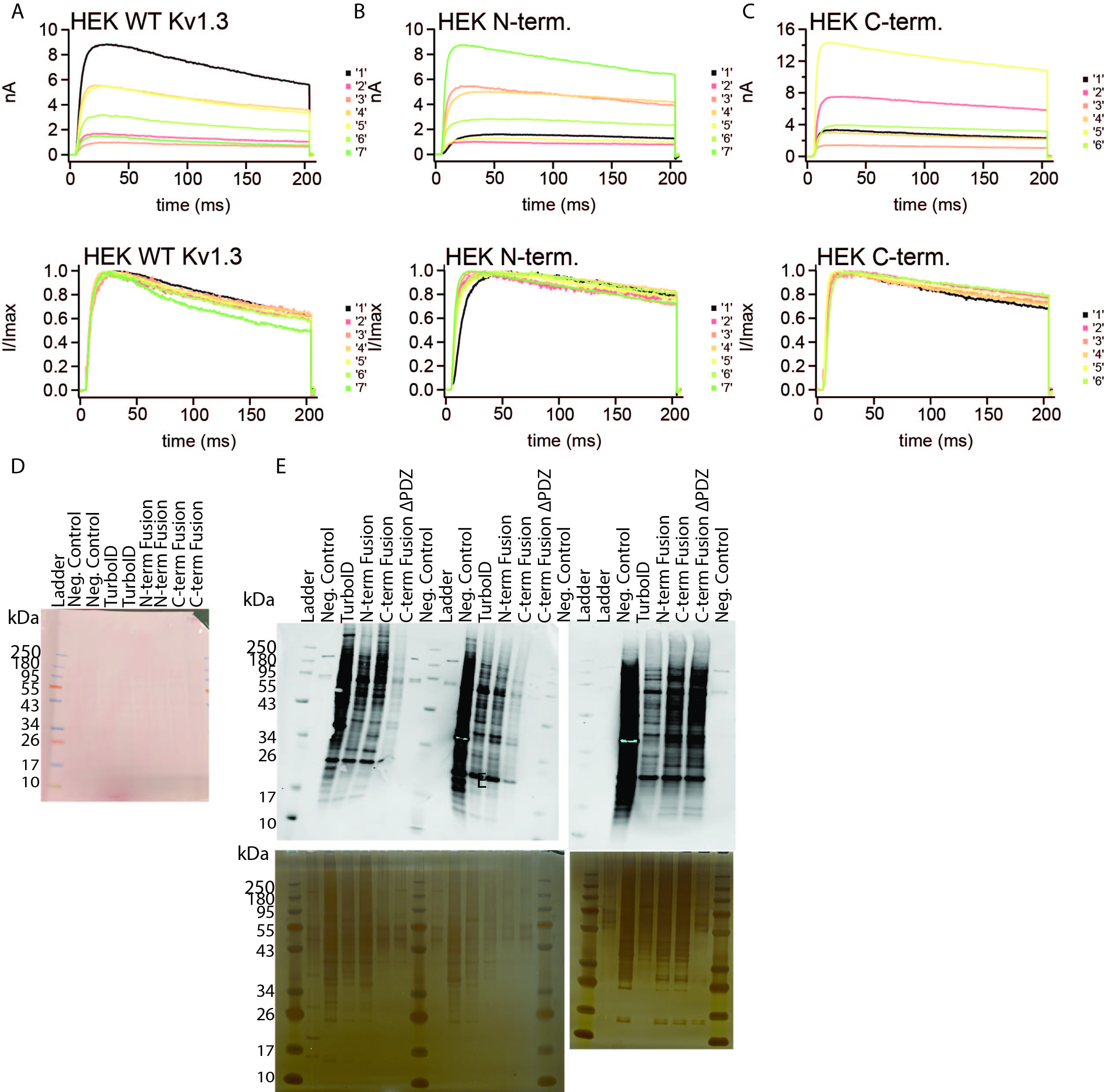

### Supplemental Figure 2

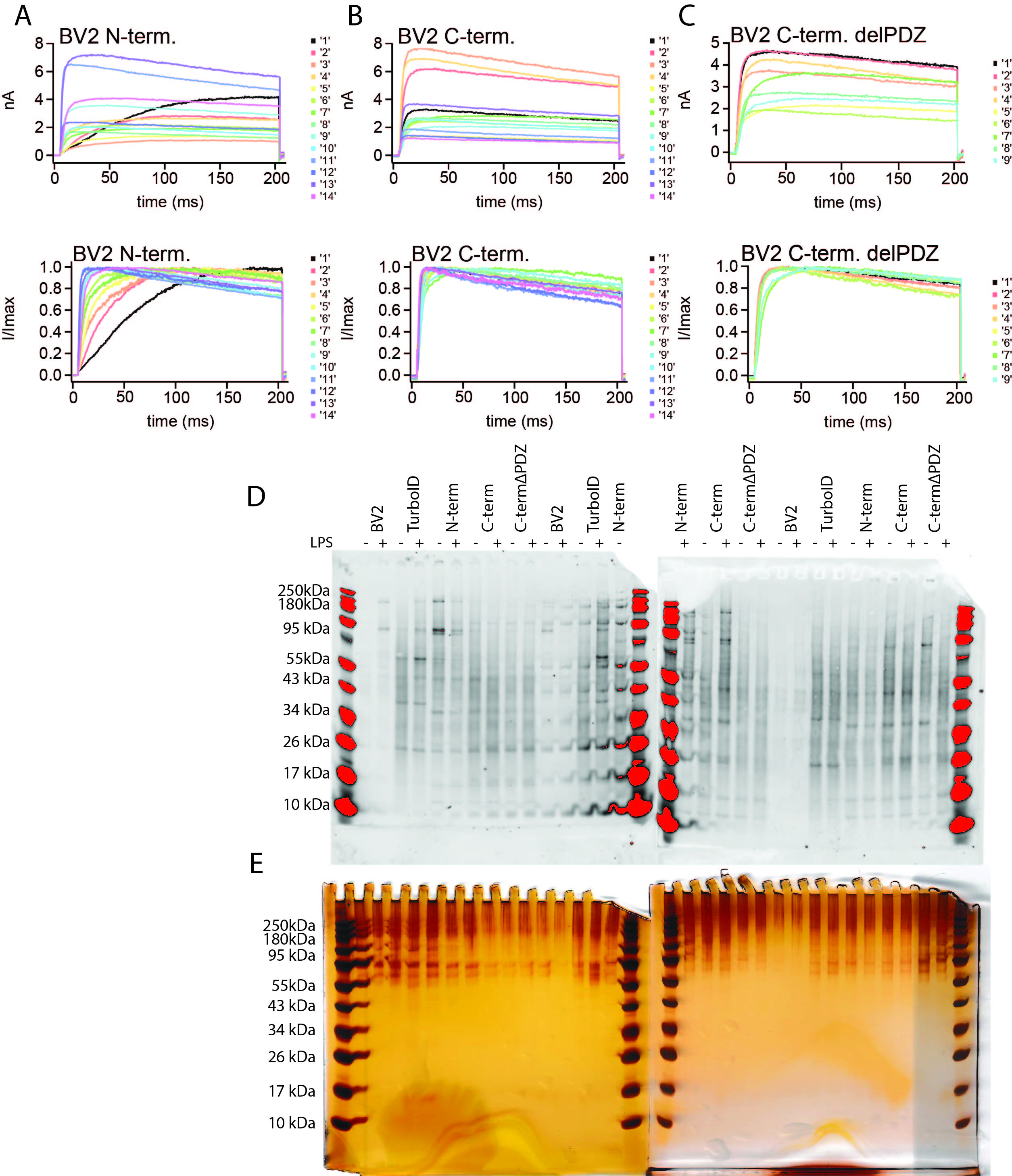

### Supplemental Figure 3

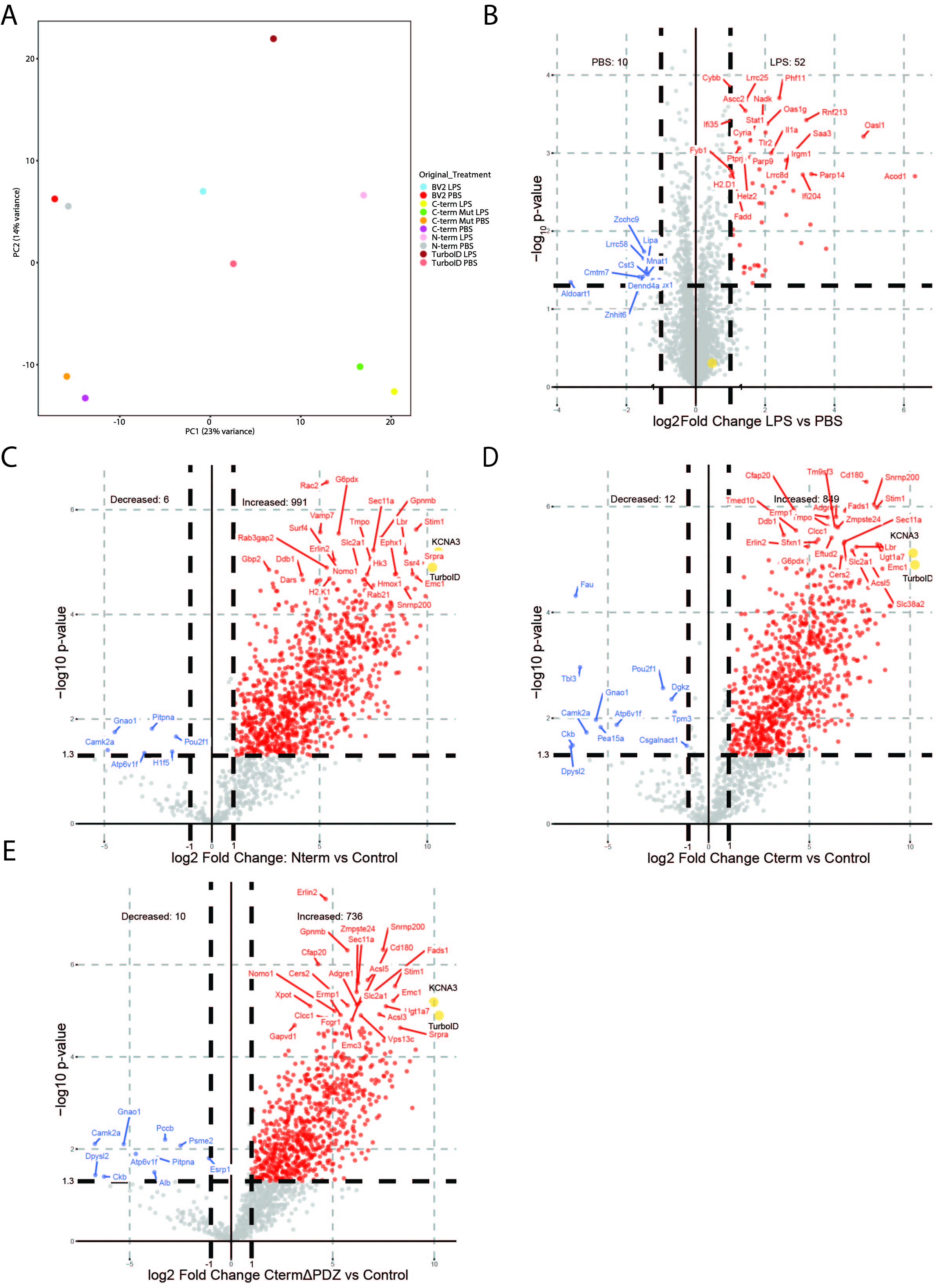

### Supplemental Figure 4

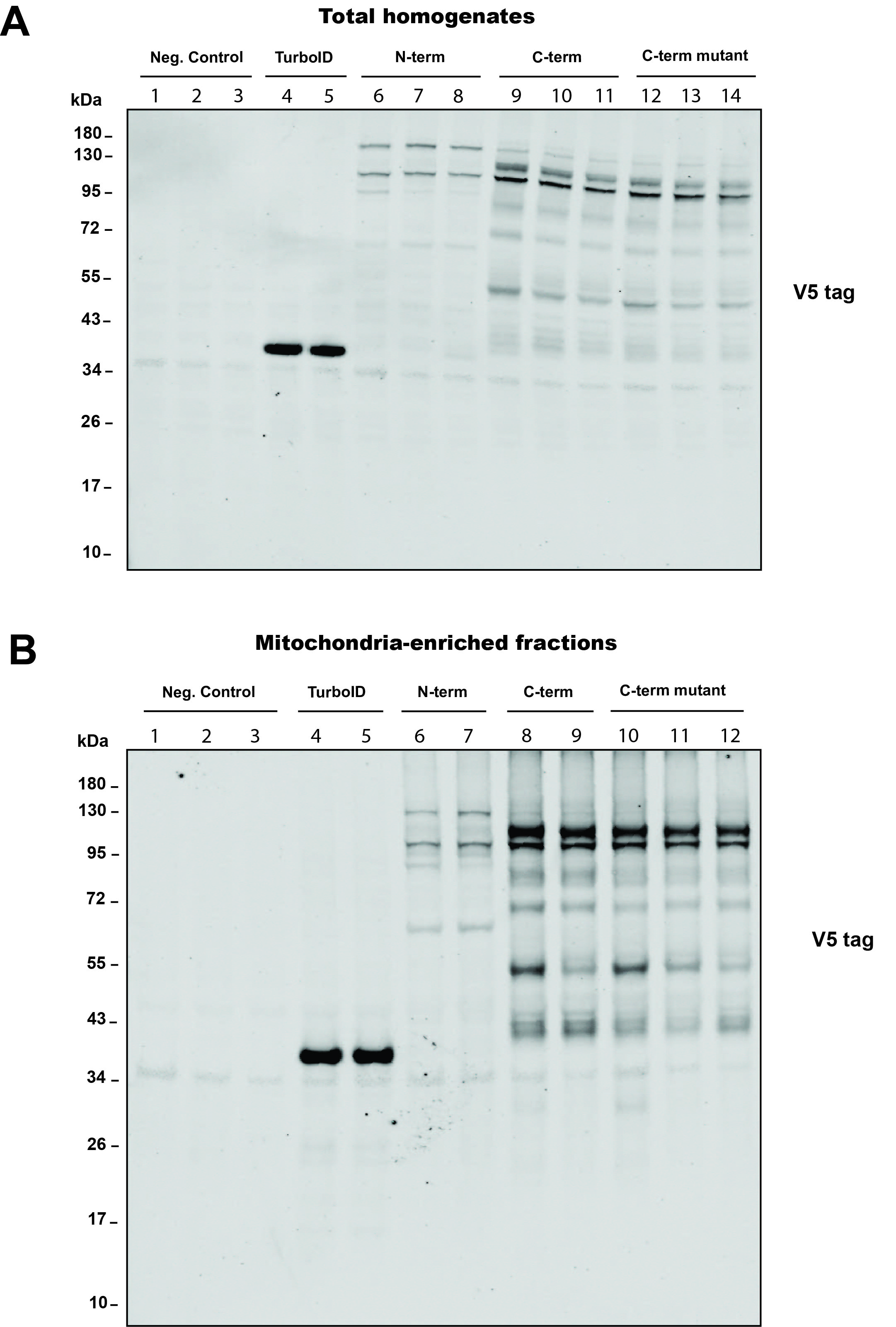
